# Competitive resource allocation drives asynchronous and rapid nuclear multiplication in the malaria parasite

**DOI:** 10.1101/2025.08.07.669072

**Authors:** Patrick Binder, Aiste Kudulyte, Severina Klaus, Thomas Höfer, Ulrich S. Schwarz, Markus Ganter, Nils B. Becker

## Abstract

The unicellular malaria parasite *Plasmodium falciparum* proliferates within the red blood cells of its human host, where it generates approximately 20 new parasites within an infection cycle of two days. Proliferation proceeds by nuclear divisions in a shared cytoplasm, followed by cellularization. In stark contrast to the highly synchronized nuclear division cycles in many eukaryotes, *Plasmodium* nuclear cycles desynchronize rapidly. Here, we elucidate the mechanism of desynchronization and study its impact on parasite fitness by combining live-cell imaging of DNA replication with biophysical modeling. We find that the standard model of independent nuclear cycles with transgenerational inheritance cannot account for the experimental data, and therefore desynchronization requires nuclear coupling. Competition for a limiting pool of a replication resource explains the data, provided that the resource is allocated sequentially to individual nuclei. Sequential allocation can be achieved by stable but reversible loading of a replication component to DNA, for example proliferating cell nuclear antigen (PCNA). Remarkably, the resultant asynchronous nuclear cycles accelerate parasite proliferation by minimizing idling times of resources. Indeed, we predict theoretically that this mechanism maximizes overall proliferation rate with limited resource availability. Thus, our findings identify nuclear cycle asynchrony as a resource-efficient means to achieve rapid proliferation.

---

Parasites are under incessant attack by the host immune system, and only a small fraction will escape host defenses and be successfully transmitted to the next host. An effective survival strategy in this situation is to compensate for the losses by rapid proliferation. This strategy is employed in particular by the eukaryotic parasites of the genus *Plasmodium* that cause the devastating disease malaria, of which *Plasmodium falciparum* is the most virulent [1]. All clinical symptoms occur during the blood stage of the infection, where *P. falciparum* proliferates inside red blood cells (RBCs), causing a parasite burden which can exceed 10 percent of infected RBCs [2, 3]. Proliferation within RBCs is achieved via the unique process of schizogony [4]. So-called merozoites invade RBCs and first develop into ring and trophozoite stages, where the parasite grows and expresses genes for host immune evasion and proliferation, remodeling the host RBC. After approx. 30 hours [5–7], the parasite proceeds to the schizont stage, which is characterized by an increasing number of individual nuclei residing in a shared cytoplasm [8]. After approx. 48 hours, cellularization occurs and around 20 daughter merozoites are released (Fig. 1a).

**FIG. 1.**
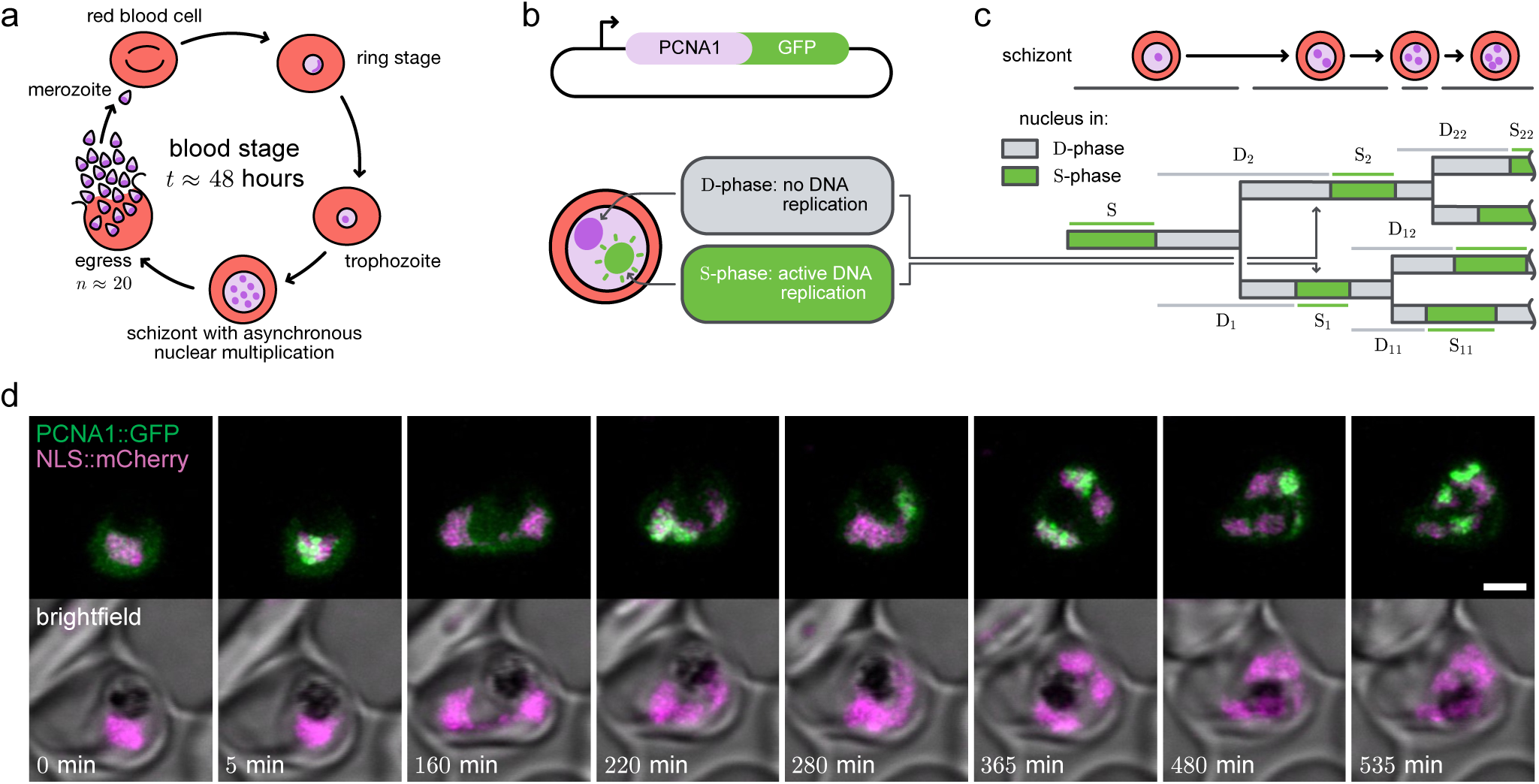
Blood-stage schizogony of the malaria-causing parasite *P. falciparum*. (a) Schematic of the asexual reproductive cycle. (b) Nuclear cycle sensor line which episomally expresses a *P. falciparum* PCNA1::GFP fusion protein. PCNA1::GFP accumulates specifically in nuclei undergoing active DNA replication, enabling live-cell tracking of S-phase nuclei. (c) Lineage tree of tracked nuclei. Nuclear cycles are divided into S-phase (active DNA replication) and D-phase (interval between to subsequent S-phases). Phase subscripts 1, 2, 12, 22 denote descendant nuclei. (d) Time-lapse confocal image sequence of a single infected red blood cell. Asynchronous DNA replications are visible from 220 min onwards. NLS, nuclear localization signal. Scale bar, 2 µm.

In marked contrast to the pattern of proliferation seen in other rapidly developing multinucleated cells, such as the early *Drosophila melanogaster* embryo [9] or the unicellular marine eukaryote *Sphaeroforma arctica* [10], nuclear multiplication in *P. falciparum* does not follow a sequence of synchronized divisions with geometric increase in number. Instead, nuclei multiply asynchronously [7, 11–13] despite sharing a common cytoplasm. In particular, the times from nuclear division to the onset of subsequent DNA replications vary considerably between sister nuclei, starting already in the first generation [13].

This asynchrony would not be possible if all nuclei were regulated by canonical eukaryotic cell cycle checkpoints [4, 14], which has been taken as indication that control of nuclear cycles is local, not global [8]. Already in 1875, botanist E. Strasburger noted that individual nuclei in multinucleated cells organize their own “sphere of influence” [15], hinting that cytoplasmic compartmentalization may be key for asynchrony. This notion is supported by data from the filamentous fungus *Ashbya gossypii*, where spatially separated nuclei show asynchrony [16, 17], but partially synchronize when brought into proximity [18–21].

By contrast, *P. falciparum* nuclei are crammed within RBCs a few µm in diameter, where any protein gradients equilibrate within milliseconds, precluding effective cytoplasmic compartmentalization. In line with this picture, homologs of the canonical G1-, Sand M-phase cyclins [22], which would then synchronize nuclei, appear to be absent [23–25]. Alternatively, asynchrony in *P. falciparum* may be due to nucleus-intrinsic mechanisms. These would give rise to stochastic nuclear cycles that are independent, except for possible correlating effects of inheritance of nuclear components such as the centrosome [8].

Here, we address the question of how and why asynchronous nuclear divisions arise in *P. falciparum*, by combining quantitative analysis of high-resolution time-lapse microscopy data with theory. Strikingly, our data rule out models where individual nuclei cycle independently as well as models where nuclei are correlated solely via inheritance of nuclear factors. This finding implies that coexisting nuclei must communicate. Specifically, the data support a biophysical model where different nuclei compete for a shared pool of diffusible (co-)factors of DNA replication, and regulation of nuclear multiplication is achieved by a limiting pool mechanism, similar to mechanisms for size regulation of cellular structures [26–29]. Thus, different from previous hypotheses [8], nuclear cycle regulation in *P. falciparum* is not nucleus-intrinsic but relies on a global, desynchronizing coupling. We also find that nuclei replicating their DNA sequentially use replication resources more steadily, which increases efficiency and speeds up nuclear multiplication in resource-limited conditions. Together, our results explain asynchrony of nuclear multiplication as a consequence of resource-efficient and fast schizogony in *P. falciparum*.

## Lineage tracking of nuclei in *P. falciparum*

To gather quantitative data on nuclear cycle timing, we used a previously generated *P. falciparum* reporter cell line [13]. This nuclear cycle sensor line ectopically expresses the red-fluorescent protein mCherry fused to three nuclear localization signals, which effectively labels all nuclei in a schizont. In addition, the line ectopically expresses the DNA sliding clamp protein PCNA1, an essential component of the eukaryotic DNA replication fork [30, 31], fused to green fluorescent protein (PCNA1::GFP). We have previously shown that PCNA1::GFP accumulates only in those nuclei of a schizont that undergo productive DNA replication (Fig. 1b) [13]. Our reporter line thus allows tracking of DNA replication and nuclear division in individual nuclei of a schizont (Fig. 1cd).

Using long-term time-lapse confocal microscopy, we had previously tracked individual nuclear lineages comprising at least the 2-nuclei stage in N = 55 cells (and one ensuing nuclear division in N = 25) [13]. To analyze these quantitative data, we divided nuclear cycles phenomenologically into two consecutive phases (Fig. 1bc). The active synthesis phase (S-phase) is defined by productive DNA replication, as reported by PCNA1::GFP accumulation in a given nucleus. The remainder of a nuclear cycle is combined into a division phase (D-phase), during which no active DNA synthesis occurs. D-phase begins when DNA replication activity terminates, extends through nuclear division and continues in each daughter nucleus until the respective ensuing S-phase begins. Thus, the S*_i_*-phase of nucleus i is followed by two D-phases D*_i_*_1_ and D*_i_*_2_ for the two daughter nuclei i1 and i2, respectively, which share a common time period from the end of DNA replication in nucleus i to its division (Fig. 1c). We chose this two-phase description (similar to earlier work [12]), because it matches the resolution of our live-cell imaging data where two phases are accurately quantifiable. By contrast, the G1, S, G2 and M phases of the canonical cell cycle are poorly defined in *P. falciparum* [4] and appear ill-suited to describe asynchronous nuclear multiplication.

## Evidence for Inter-Nuclear Coupling

We asked if individual nuclei multiply autonomously, that is, without physical coupling to other nuclei coexisting in the same cell. In the simplest such model (model 1) cycle regulation relies purely on non-heritable nucleus-intrinsic factors and no signaling occurs between nuclei. All nuclei are equally DNA-replication competent. Mathematically, these requirements lead to a two-phase branching process [32, 33], where each S- or D-phase is drawn independently from the corresponding empirical phase distribution, derived from experimental data of the first two generations.

To test model 1, we measured the S-phase durations of sister nuclei at the 2-nuclei stage. While model 1 predicts no correlation between the S-phase durations of autonomous nuclei, the data exhibited a positive correlation between sisters, especially when S-phases were longer (Fig. 2a). Because model 1 failed to capture the correlation, we ruled it out.

**FIG. 2.**
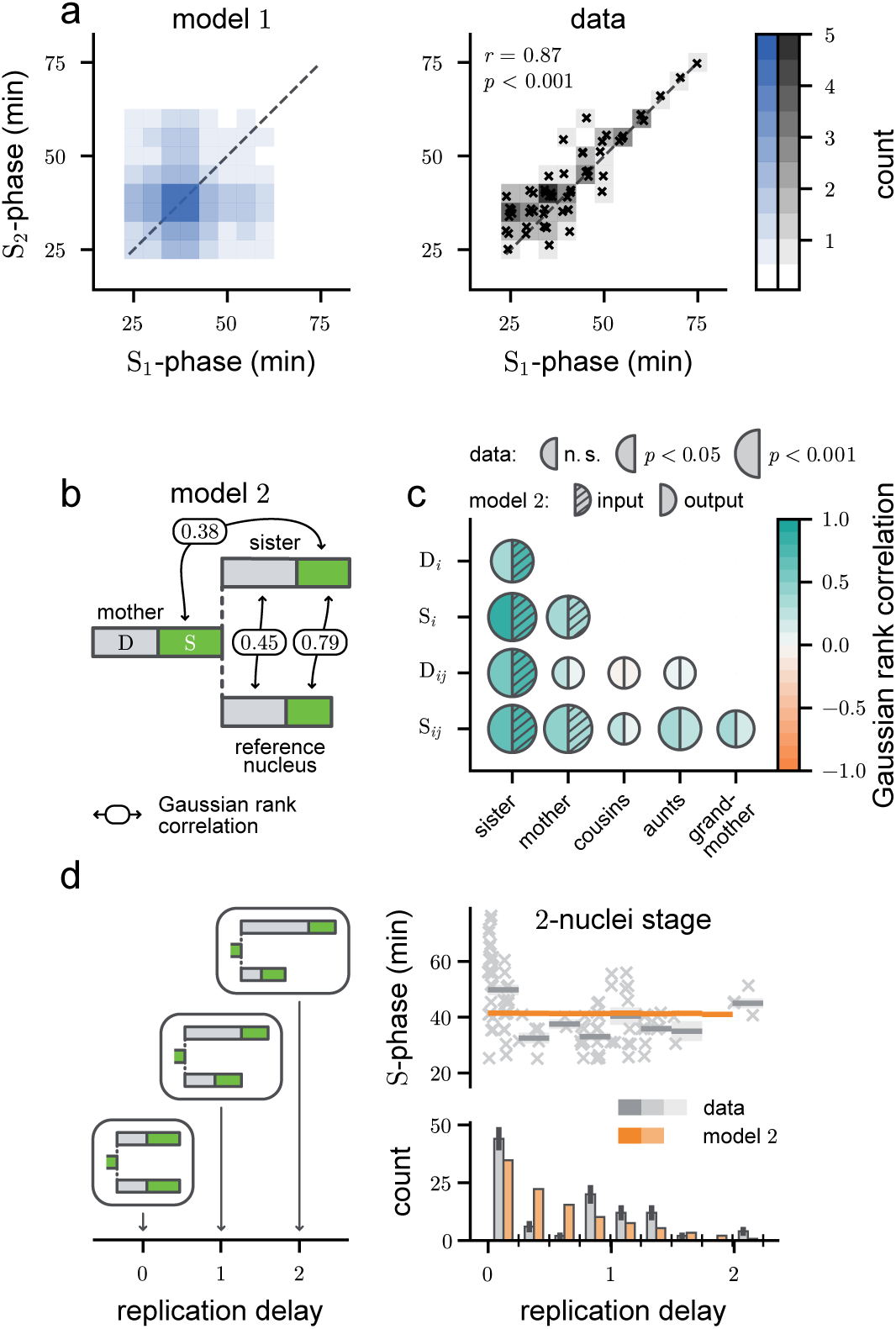
Observed and modeled nuclear cycle correlations in *P. falciparum*. (a) The two-phase branching process (model 1, left) fails to reproduce the positive correlations observed (right) between sister S-phases at the 2-nuclei stage. *r*: moment correlation coefficient with *p*-value, *N* = 55. (b) Schematic of model 2, which incorporates inheritance from mother to daughter nuclei, with included Gaussian rank correlations. (c) Correlation structure. Disks: data (left, with *p*-value) and model 2 (right; hatched: input, empty: prediction). (d) S-phases vs. delay at the 2-nuclei stage. Left: Illustration of replication delay, defined as (*τ*_D2_ − *τ*_D1_)*/*⟨*τ*_S1_ ⟩, where S_1_ is the sister S-phase that starts first. Top right: S-phase duration versus delay. Gray crosses, lines and shading: data, mean, and its bootstrapped standard error, respectively; *N* = 102. Orange lines: model 2 prediction. Bottom right: delay histogram with bootstrapped standard error bars.

Next, we included inheritance of nuclear factors from the mother as a natural mechanism that would correlate sister S-phases. To this end we constructed model 2 as a Markovian bifurcating autoregressive (BAR) process [34, 35]. Nuclei in model 2 are uncoupled, meaning that their cycle phases are not directly affected by coexisting nuclei. Inheritance is incorporated by drawing the duration of nuclear cycles phases in sister nuclei from distributions that may depend on the phases realized in the mother. For instance, both sister nuclei can inherit a bias for a longer S-phase from a mother with a long S-phase, which would correlate sister S-phases positively. Furthermore, to capture the fact that sister D-phases have a common initial period (from the end of the maternal S-phase to division), we also allow for direct sister phase correlations in the model. More precisely, we analyzed *P. falciparum* schizonts with 1 to 4 nuclei for the Gaussian rank correlations between nuclear phases. The joint distribution of the six phases of mother and daughters in model 2 was then parameterized by including all significant correlations, setting insignificant correlations to zero (Fig. 2bc; see Fig. S1 and SI Sec. I for details). For instance, S-were uncorrelated with D-phases (Fig. S1c), while S-phases were positively correlated between mother and each daughter.

By construction, model 2 reproduced the single-phase distributions and mother-daughter and sister correlations present in the data (Fig. 2c). Model 2 then also successfully predicted observed correlations between more distantly related nuclei. For instance, S-phases of aunts and nieces were positively correlated.

To further test model 2, we focused on the prolonged S-phases of sister nuclei that commence DNA replication simultaneously, which we had found previously [13]. We first quantified the delay between two sister nuclei entering S-phase. We define the replication delay (D_2_ − D_1_)/⟨S_1_⟩ as the difference of replication start times, normalized by the average duration of the first (leading) S-phase. Thus, a replication delay of 0 indicates simultaneous entry into S-phase, a delay of 1 implies roughly sequential DNA replication, and a delay > 1 indicates a gap between the end of S_1_ and the onset of S_2_ (Fig. 2c, left panel). While our experimental data showed an almost two-fold slowdown of S-phases in sister nuclei with replication delay 0 (Fig. 2d, top right panel, gray points and lines), model 2 predicts a constant S-phase duration at all replication delays (orange lines).

We next also compared the frequencies of the various replication delays (Fig. 2d, bottom left panel). While model 2 predicts a gradual decrease of events with increasing delay, our experimental data showed a bimodal distribution with enriched replication delays 0 and 1 and depleted delays around 0.5.

Because both the prolonged S-phases at replication delay 0 and the bimodal distribution of delays were not captured by model 2, we ruled it out as explanation for the observed behavior. We also excluded extended BAR models that incorporate more distant correlations, as these would not correspond to physiological mother-daughter inheritance.

In conclusion, our experimental data rule out a large class of models in which the duration of the different phases of nuclear multiplication are determined purely by inheritance of nucleus-intrinsic factors, but not by other nuclei coexisting in the cell. This strongly suggests that some physical mechanism couples the nuclei in the common cytoplasm.

## A limiting pool mechanism couples replicating nuclei

A simple and parsimonious mechanism that would couple replicating nuclei is competition for shared resources needed for DNA replication, *i.e.*, components (cofactors) of the replication machinery.

Competition for resources is central for so-called limiting pool mechanisms, which can regulate the size of cellular structures and organelles without the need for dedicated size sensing pathways [26, 27]. The basic feature is that as organelles grow, they incorporate subunits from a cytosolic pool, depleting it, and slowing further growth. This mechanism has been shown to effectively regulate the size, the scaling relative to cell size and the scaling among a set of competitor organelles, such as mitochondria, vacuoles, nuclei, centrosomes and flagella [29], mitotic spindles [36], and actin networks [28].

Similar to sharing of (or competition for) cellular building blocks, nuclei in a *P. falciparum* schizont share replication cofactors: For instance, episomally expressed PCNA1::GFP translocates between nuclei in S-phase within minutes while conserving the total amount in the cell [13], e.g., the mammalian homolog PCNA has been shown to be loaded onto replication forks reversibly [37]. Thus, in analogy to the size regulation by limiting pool mechanisms, we hypothesized that there exists a (yet unspecified) shared cofactor R, called ‘resource’ below, which limits the speed of DNA replication. If R was distributed unevenly, nuclei with little R would not replicate their DNA, which can cause the observed asynchrony. Alternatively, if R was distributed evenly but in insufficient amounts, simultaneous S-phases would be prolonged.

At variance to the limiting pool mechanisms mentioned above, the cofactor R is not permanently incorporated into nuclei, but released back to the cytoplasm after S-phase. Although not successively depleted, in a growing population of nuclei the initial R pool would nevertheless soon become a bottleneck for DNA replication, necessitating some form of resupply.

To explore the implications of a limiting pool of resource quantitatively, we constructed model 3, which couples nuclei via simple resource kinetics (Fig. 3a). Model 3 posits that nuclei exist in the replication-competent S^∗^-phase, or in D^∗^-phase. (These two phases differ from the experimentally observed phases S and D through a threshold in intensity that has to be exceeded for experimental observation, see below.) Free replication resource R translocates between nuclei and the cytoplasm, equilibrating rapidly within the entire *P. falciparum* cell (Fig. 3b). In any S^∗^-phase nucleus i, resource R reversibly associates with the DNA replication forks F*_i_*, forming the active replication fork complex:

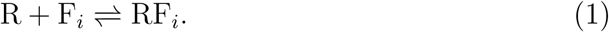

**FIG. 3.**
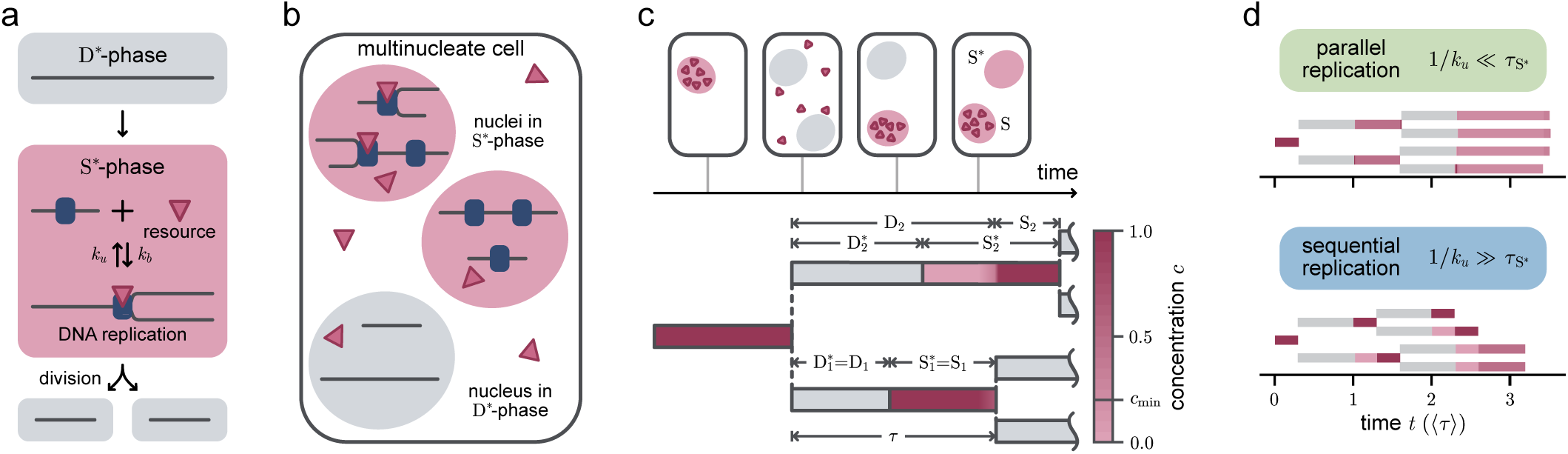
Biophysical model for competitive resource sharing (model 3). (a) In D^∗^-phase, the replication machinery is disassembled. Upon entering S^∗^-phase, the replication machinery is assembled, but active replication in addition requires loading of replication resource molecules. (b) S^∗^-phase nuclei within a schizont compete for replication resource in the cytoplasm. (c) After the lagging daughter nucleus (subscript 2, top) enters S^∗^-phase, it queues for resource to become available, remaining assigned to D-phase. It enters productive DNA replication (S-phase) after the leading daughter is done. (d) Example lineage trees. Parameters: *k_u_* = 0.01 (sequential replication) and *k_u_*= 100 (parallel replication); *r*(*t*) ≡ 1, ⟨*τ* ⟩ = 1, *τ*_S_∗ = 0.3, *σ*_D_∗ = 0.02, *k_b_* = 10^6^.

In turn, active forks RF*_i_* replicate DNA, so that the DNA amount increases from 1 to 2 haploid genomes during each S^∗^-phase, without resource consumption.

Whenever the fraction of activated forks in nucleus i exceeds a threshold, it is experimentally detectable by localization of R in nucleus i, which is then classified as being in active S-phase, contained within S^∗^-phase. Indeed, such a scenario may be realized for PCNA1::GFP, which transiently accumulates in those nuclei that replicate their DNA [13]. However, model 3 is agnostic about the molecular identity of R.

S^∗^-phase ends when a nucleus has fully replicated its DNA, at which point replication forks disassemble, and the bound resource is instantaneously and irreversibly released. Two stochastic D^∗^-phases ensue, which span nuclear division and end at the start of S^∗^-phase in the two daughter nuclei, respectively. As schizogony proceeds, both randomness in D^∗^-phase durations and differences in DNA replication speeds contribute to the desynchronization of nuclear cycles.

We now give a full account of model 3, completing the qualitative description above by defining equations. In *P. falciparum*, licensed origins of replication are likely abundant [5, 38], and the overall DNA replication rate in a nucleus is limited by the number of replication forks (initiated at licensed and activated origins). For simplicity, we consider the total number of replication forks f*_i_* in any S^∗^-phase nucleus to be identical, and we measure all molecule numbers relative to f*_i_*. Then in any S^∗^-phase nucleus i we have 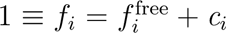, where 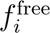 is the amount of inactive forks F*_i_* without loaded resource, and *c_i_*, the amount of active resource-fork complexes RF*_i_*, respectively. In D^∗^-phase nuclei no forks exist, and 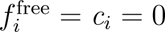 (Fig. 3ab). Summing over all nuclei, 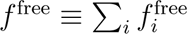 and *c* ≡ ∑*_ii_ c_i_*, respectively, we then find that f^free^ + *c* = n_S_∗, the number of nuclei in S^∗^-phase at any given time. We also introduce the amounts of free resource r^free^ and total resource r ≡ r^free^ + *c*. In our units, r also equals the maximum number of nuclei that could be fully engaged in active DNA replication at any given time, in the case that r = c and no free resource remains, r^free^ = 0.

With these definitions, mass-action kinetics for resource loading and unloading in nucleus i (Eq. 1) read

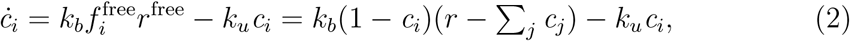

a system of coupled ordinary differential equations for the set of active forks {*c_i_*}, driven by the time-varying total resource r. The coupling arises via the shared pool r^free^ of free resource for which S^∗^-phase nuclei compete (Fig. 3c). Importantly, the inverse rate constant 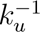 sets the lifetime of active forks. The timescale for fork activation is set by (rk*_b_*)^−1^ in the resource-rich regime.

The DNA synthesis rate *g⠁_i_* in nucleus *i* is taken to be proportional to the number *c_i_* of activated forks:

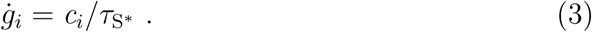

This makes the assumption that replication forks do not interact, e.g., by competing for nucleotides [39]. The DNA amount g*_i_* increases from 1 to 2 haploid genomes during S^∗^-phase. Although the time point of nuclear division is not resolved in the model, for simplicity, g*_i_* is reset to 1 at the start of each daughter D^∗^-phase. The total DNA amount *g* ≡ ∑*_i_ g_i_* increases continuously with time [6, 40]. A fully activated nucleus (*c_i_* = 1) is seen to replicate at a maximal rate 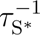, so that *τ_S_*∗ can be recognized as the minimum possible S-phase duration in the model.

Finally, motivated by the shape of observed D-phase durations, D^∗^-phases are modeled as Gamma-distributed, parameterized by the mean ⟨τ_D_∗ ⟩ and standard deviation σ_D_∗ . In resource-rich conditions, the cycle time is given by

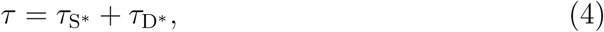

and it follows a Gamma distribution shifted by *τ_S_*∗, with mean ⟨*τ*⟩ = *τ_S_*∗ + ⟨*τ_D_*∗ ⟩ and standard deviation *σ_D_*∗ . In resource-limited conditions, the cycle time is increased by slowed DNA replication.

To relate this model to data, we account for the fact that active DNA replication in a given nucleus is detectable only above a threshold size [13, 40]. Concretely, we assign nucleus i to S-phase when activated forks exceed a detection threshold *c_i_* > *c*_min_ ≈ 20%. Notably, detectable S-phases can begin well after S^∗^-phase if a lack of available resource slows initial replication. Nuclei not in S-phase are assigned to D-phase. Thus, each S^∗^-phase contains a detected S-phase, while each D-phase contains a D^∗^-phase (Fig. 3c).

## Resource loading kinetics and abundance control the synchrony of nuclear multiplication

Next, we investigate how synchronous or asynchronous nuclear cycles arise in model 3. Motivated by the observed kinetics of PCNA1::GFP [13, 40] we focus on a regime characterized by high resource-fork affinity, k_u_/*k_b_* ≪ 1, and rapid binding, *k_b_τ_S_*∗ ≫ 1. It turns out that the amount of available resource in relation to replicating nuclei is a key determinant of the nuclear cycling dynamics. We call a cell resource-limited whenever n_S_∗ > *r*, which implies that one or more of the S^∗^-phase nuclei replicate at a reduced rate 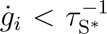 due to lack of resource. In resource-limited cells, due to high affinity, essentially all available resource is bound to replication forks, leaving no free resource unbound (see SI Sec. II).

Now consider a nucleus i as it enters S^∗^-phase. In the resource-limited cell, R is already fully in use by other running S-phases. How quickly nucleus i can begin DNA synthesis, depends on the kinetics of R release from other nuclei. Fast release, 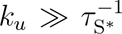, equilibrates R quickly among all nuclei, enabling immediate DNA replication in nucleus i while slowing other S-phases. Thus, fast resource release entails parallel DNA replication in S^∗^-phase nuclei. Conversely, slow release, 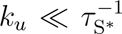, lets already-DNA replicating nuclei retain all loaded resource. S-phase in nucleus i is delayed until resource R is freed at the end of some other S^∗^-phase. Hence, slow release leads to sequential DNA replications.

In sum, by adjusting *k_u_*, model 3 interpolates between parallel and sequential modes of DNA replication (Fig. 3d). Resource shortage results in prolonged, synchronized S-phases, or asynchronous S-phases, respectively. Abundance of resource will enable fast, immediate and synchronized DNA replication in either mode, hiding release kinetics.

As time passes and nuclei increase in number, any fixed resource pool r will soon limit nuclei to linear growth, as the overall rate of DNA synthesis in the cell is bounded by r/τ_S_∗ (cf. Eq. 3). However, *P. falciparum* exhibits exponential nuclear multiplication and DNA synthesis over several generations (Fig. S2, [6, 13, 41]). To complete model 3, we thus need to specify an increasing total available resource *r* = *r*(*t*). In particular, exponential nuclear multiplication requires that resource also increase exponentially, which we incorporate in an effective way by setting

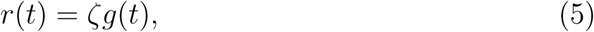

where the resource availability ζ is a parameter that quantifies the degree to which synthesis of resource proteins keeps up as schizogony proceeds. In particular, we find that when ζ exceeds some critical value ζ*_c_*, resource is not limiting (in fact we can show ζ*_c_* < 1, see below and SI Sec. III).

## Nuclear multiplication in exponentially growing populations

Next, we aim to characterize the dynamics of nuclear growth in model 3, for which it is convenient to consider the total DNA synthesis rate *g*⠁ = *c*/τ_S_∗ (summing Eq. 3 over all nuclei). To relate the synthesis rate to resource production, we introduce the instantaneous resource utilization parameter

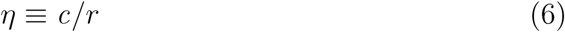

as the instantaneous fraction of available resource used for DNA replication. Intuitively, η ≪ 1 indicates unused potential for growth, and *η* = 1 corresponds to maximal resource usage. Combining with Eq. 5 we obtain

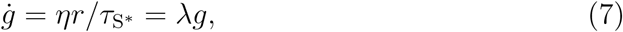

where

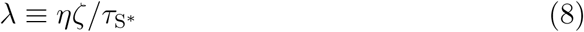

is the instantaneous exponential growth rate. Eqs. 7–8 express DNA replication in terms of the intrinsic synthesis rate 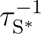, external control by resource availability ζ, and resource utilization η resulting from cycle phase timing.

We first focus on nuclear populations in the long-time limit. As time progresses, randomness in the D^∗^-phase durations results in gradual desynchronization of nuclear multiplication. Eventually, Eqs. 2–7 attain a state of steady growth, which is characterized by exponentially increasing DNA content, total resource and nuclear number, but stationary population structure (Fig. 4a).

**FIG. 4.**
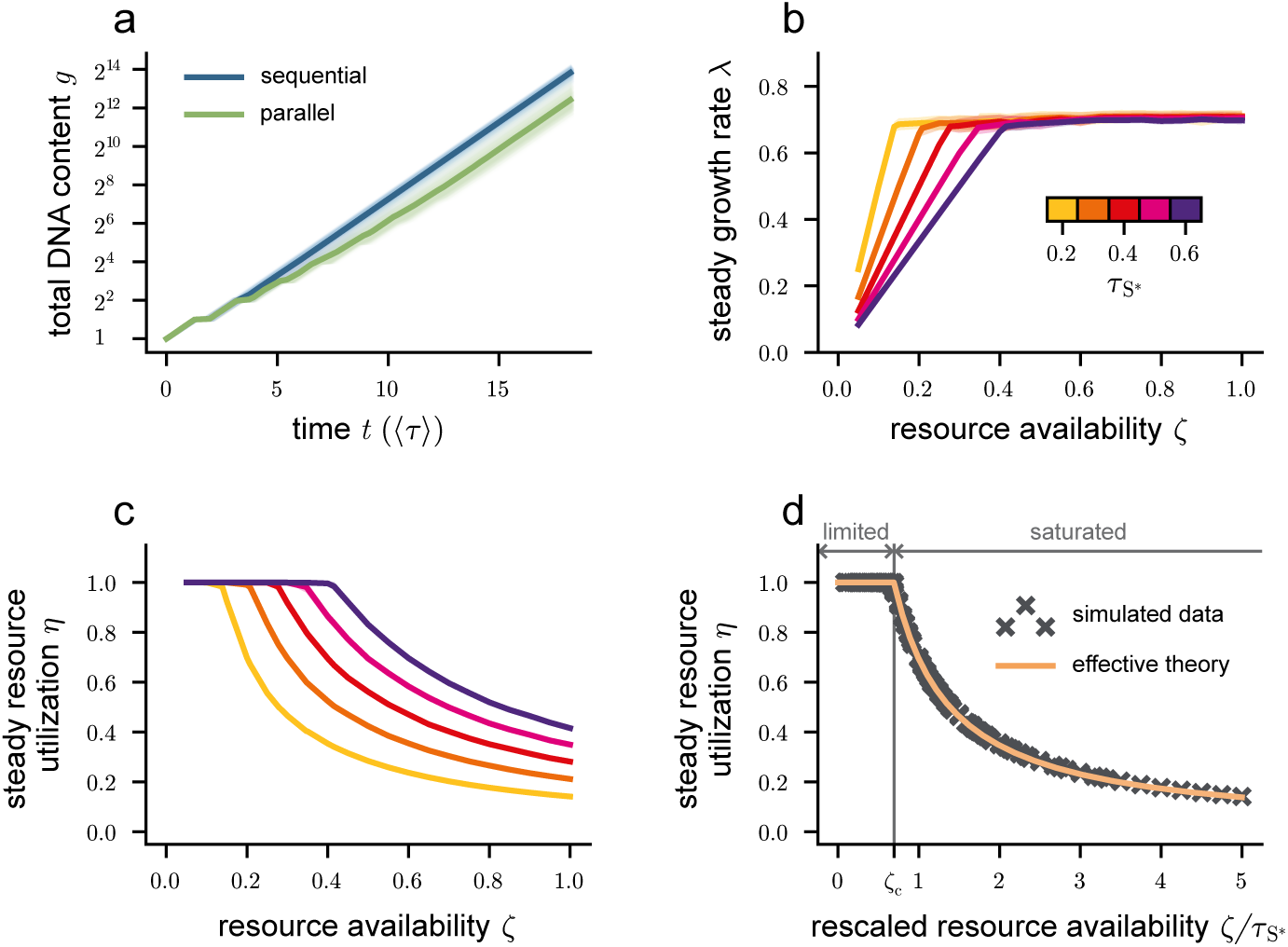
Asymptotic regime of nuclear multiplication in model 3. (a) Time course of total DNA content *g*(*t*) showing that convergence towards steady exponential growth is slower in parallel mode than in sequential mode. Lines, shading: simulation mean and single realizations. (b) Steady growth rates *λ* vs. resource availability *ζ*. Growth rates saturate to *λ*_max_ at the critical *ζ_c_* = *λ*_max_*τ*_S_∗ . (c) Steady resource utilization *η* vs. *ζ*. Utilization decreases proportionally to 1*/ζ* above *ζ_c_*. In b and c, sequential and parallel mode curves are identical. (d) Collapse of simulation results onto the theoretical prediction given by Eq. 8 (yellow line). Crosses represent a simulation with a specific set of parameters (*τ*_S_∗ *, ζ, k_u_*). In a-d, 480 lineage trees containing approximately 3 · 10^4^ S^∗^-phases were generated per parameter set. The growth rate *λ* was determined by fitting an exponential to *g*(*t*) over the last 3 generations; *η* was obtained as *c/r* at the end of the simulation. Parameters: ⟨*τ* ⟩ = *τ*_S_∗ + ⟨*τ*_D_∗ ⟩ ≡ 1 (fixed), *σ*_D_∗ = 0.1, *k_b_* = 10^6^, *k_u_* = 0.01 (sequential mode) and *k_u_* = 100 (parallel mode). In panel a, *ζ* = 0.17 *< ζ_c_* and *τ*_S_∗ = 0.3.

To determine the steady-growth values of η and λ, first consider resource limited cells (i.e., r < n_S_∗). High RF binding affinity ensures that resource is fully bound to forks at all times, i.e., r^free^ = 0 and therefore η = 1. Eq. 8 takes on the constant value λ = ζ/τ_S_∗ .

Conversely, in non-resource limited cells where r > n_S_∗, resource fully saturates available forks. Thus, every S^∗^-phase nucleus i has *c_i_* = 1 and completes S^∗^-phase after the minimal duration τ_S_∗ ; the full nuclear cycle takes τ = τ_S_∗ + τ_D_∗ (Eq. 4) where only τ_D_∗ is stochastic. As a result, the population growth rate attains its maximal value λ_max_, given implicitly by the Malthusian relation (e.g. [32], see also SI Sec. IV)

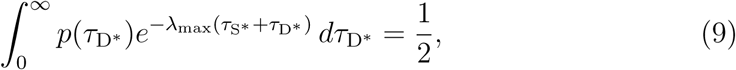

from which η = λ_max_τ_S_∗ /ζ can be obtained by Eq. 8.

Combining the cases, we obtain the following picture: As the resource availability ζ is increased, the steady growth rate first increases linearly and then saturates to λ_max_ at a critical ζ*_c_* = λ_max_τ_S_∗ (Fig. 4b). In turn, the resource utilization remains at η = 1 up to ζ*_c_* and then decreases as η ∝ 1/ζ (Fig. 4c). These relations can be summarized as

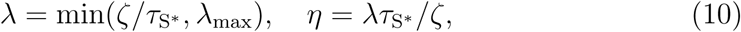

with λ_max_ given by Eq. 9. In the limit of deterministic τ_D_∗, we can solve Eq. 9 explicitly to obtain

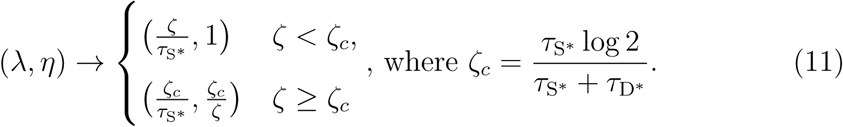

Interestingly, the resource release kinetics, which control parallel vs. sequential S^∗^-phases in the initial generations, have no bearing on the eventual steady growth rate, as k*_u_* does not appear in Eq. 10. Thus, any difference in growth rate due the amount of transient initial synchronization disappears once nuclei are fully desynchronized (Fig. 4a, and 4bc where sequential and parallel mode simulations give identical results).

By contrast, the intrinsic balance between the nuclear cycle phases has an effect: In resource limited cells, decreasing the S -phase fraction τ_S_∗ /⟨τ⟩ increases the steady growth rate (Fig. 4b). This is because a shorter S^∗^-phase reduces the fraction of nuclei that require resource at any given time, effectively increasing availability of the resource (cf. Eq. 8 with η → 1). In absence of resource limitation (beyond ζ*_c_*), timely resource release is not needed and growth saturates at λ_max_, independent of S -phase fraction. Indeed, Eq. 9 is invariant under shifts of p(τ_D_∗) and τ_S_∗ by equal and opposite amounts, which do not change τ .

In Fig. 4d, simulation data across a wide range of the parameters τ_S_∗ /⟨τ⟩, σ*_D_* and ζ, spanning resource limited and non-limited cells, collapse onto a master curve. The collapse confirms our effective theory for steady growth and its central requirement of full desynchronization of nuclear cycles in the long-time limit.

Next, we utilized model 3 to investigate the initial transient growth phase. Nuclear multiplication begins with a single (trivially synchronized) nucleus, and then gradually desynchronizes during several rounds of nuclear multiplication (Fig. 5a). The crucial feature that distinguishes partially synchronized from fully asynchronous populations is the existence of time periods where most of the nuclei are in D^∗^-phase, which causes an intermittent drop in resource utilization η and DNA synthesis ġ. Such gaps in resource utilization are more prominent and persistent in populations of nuclei that replicate their DNA in parallel, and where cycles thus remain synchronized much longer (Fig. 5ab). To quantify the impact of gaps in resource utilization on nuclear multiplication, we obtain from Eq. 7 the ratio between total DNA content from sequential vs. parallel replication mode

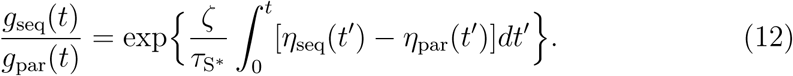

**FIG. 5.**
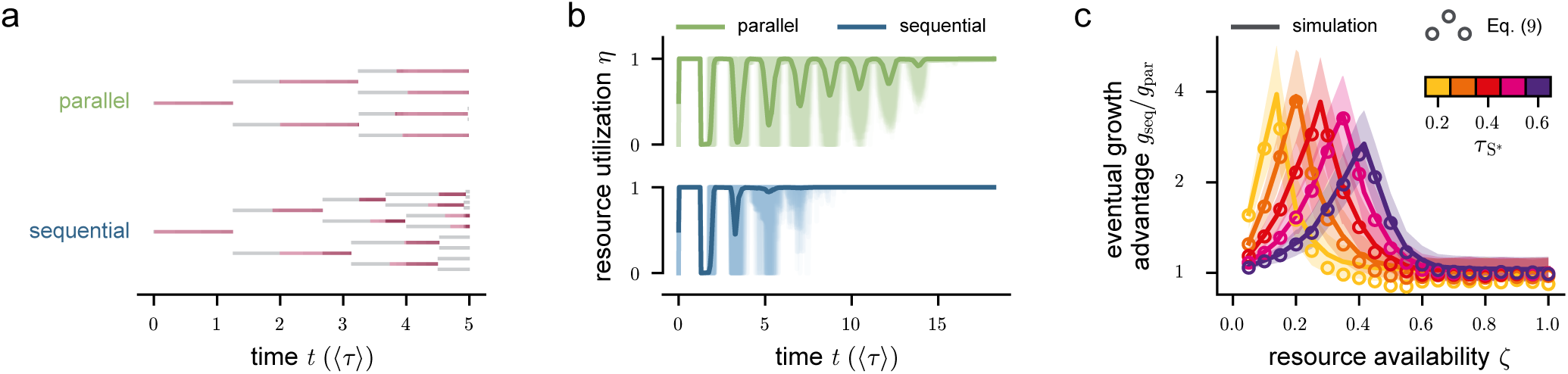
Initial phase of nuclear multiplication in model 3, highlighting the impact of replication mode on resource utilization and growth. (a) Example lineage trees for parallel and sequential replication modes. (b) Time course of resource utilization *η*. Transient drops in *η* (gaps) due to synchronized D^∗^-phases persist longer in parallel mode. Lines, shading: simulation mean and single realizations. (c) The eventual growth advantage of sequential mode is maximized at intermediate resource availability. Lines, shading: simulation mean and standard errors of *g*_seq_*/g*_par_ at the end of the simulation, respectively. Circles: prediction by Eq. 12. Parameters as in Fig. 4, and in a, b as in Fig. 4a.

Eq. 12 shows that the time integrated resource utilization controls the speed advantage afforded by the choice of replication mode.

In simulations, we find that the sequential DNA replication mode has a net utilization advantage η_seq_ − η_par_ ≥ 0 during the first few nuclear cycles. Importantly, as predicted by Eq. 12, the initial utilization advantage entails a persistent growth advantage, even though η_seq_ − η_par_ → 0 in the long term (Fig. 5c). The extent of the advantage depends on the resource availability. Deep in the resource saturated regime ζ ≫ ζ*_c_*, nuclei replicate at maximal rate during the entire transient desynchronization period, leaving no advantage. At low ζ ≪ ζ*_c_*, the advantage also disappears, as the growth rate λ is of order O(ζ), Eq. 8. At intermediate availability, the sequential mode has a sizable advantage which can reach up to threefold for plausible values τ_S_∗ ≃ 0.5, ζ ≃ 0.3, and σ_D_∗ ≃ 0.1 (Fig. 5c). More generally, sequential-mode DNA replications accumulate an advantage for as long as the parallel-mode population is slowed by the gaps in resource utilization (Fig. 5b). Further reducing σ_D_∗ can therefore further increase the growth advantage, limited only by inevitable baseline fluctuations in the duration of D^∗^-phases.

In sum, the sequential mode of DNA replication enables faster nuclear multiplication under a resource constraint. It does so by queuing replication-competent S^∗^-phase nuclei for DNA replication during the first few generations. Queuing delays the onset of DNA replication in some nuclei, but it speeds DNA replication in replicating nuclei, allowing them to proceed to D^∗^-phase earlier. Resource freed early has a high probability of being reused immediately in a queued nucleus. Thus, queuing leads to more efficient resource usage and ultimately, to a persistent growth advantage.

## A limiting pool mechanism can explain asynchronous nuclear multiplication in *P. falciparum*

As shown above, sequential DNA replication is a general strategy to accelerate the initial rounds of nuclear replication with limited resources. Our imaging data show that nuclei in *P. falciparum* desynchronize early (Fig. 2d and S2b) [13]. Is a limiting pool mechanism indeed responsible for this desynchronization, and does it have speed benefits? To address these questions, we simulated nuclear multiplication aiming to reproduce essential features of the data, in particular, the prolonged S-phase of simultaneous sister nuclei, and the bimodal distribution of replication delays.

To match experimental data, we extended model 3 in three ways. First, because we previously found that one round of *P. falciparum* schizogony produces n ≃ 24 daughter nuclei [13], we prohibit further entries into S^∗^-phase after 23 S^∗^-phases have been initiated, implementing a counter mechanism as described previously [13, 41].

Second, our initial simulations showed that the simple exponential resource model Eq. 5 supplies too much available resource towards the end of schizogony. Consistently, we had found previously that although nuclei initially multiply exponentially (Fig. S2), the multiplication rate slows down as the cell approaches the end of schizogony [13]. To account for the slowdown, we replaced Eq. 5 by a more realistic model, where the total amount of resource r(t) mirrors the measured time course of ectopically expressed PCNA1::GFP in *P. falciparum* [13], allowing for a constant scaling parameter ξ defined such that r(t) → ξ as t → ∞. In this setting, r(t) grows initially but saturates approximately halfway through the schizont stage, i.e., after 2–3 rounds of multiplication (Fig. S3a).

Third, there is considerable variability in overall durations of nuclear multiplication across different parasites [7, 13, 41], which we incorporated by sampling ξ per parasite from a uniform distribution with adjustable range. Details of the model extension and the qualitative fit procedure are given in SI Sec. VI.

While the uncoupled models 1 and 2 had failed to match our data on the 2-nuclei state (Fig. 2), model 3 was able to reproduce both the prolonged S-phase of simultaneously DNA replicating nuclei and the bimodal distribution of replication delays (Fig. 6a). Critically, this match was only achievable with slow resource release (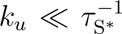), i.e., in sequential replication mode, but not in parallel mode (Fig. S4). To rationalize the underlying mechanism, consider a resource-limited cell in sequential mode at the 2-nuclei stage. The lagging sister nucleus will usually be queued for replication until the leading sister finishes S^∗^-phase; this generates the mode at delay 1 of the bimodal S-phase distribution. Still, simultaneous S^∗^-phases can arise if both sisters enter S^∗^-phase with a delay shorter than the (fast but finite) resource binding timescale (k*_b_*r)^−1^, before the leading sister has had time to sequester all resource. These events generate the mode at short delays ≃ 0. Only the simultaneous S-phases have to share the limited resource and only these are therefore slowed (Fig. 6b), as observed in the data.

**FIG. 6.**
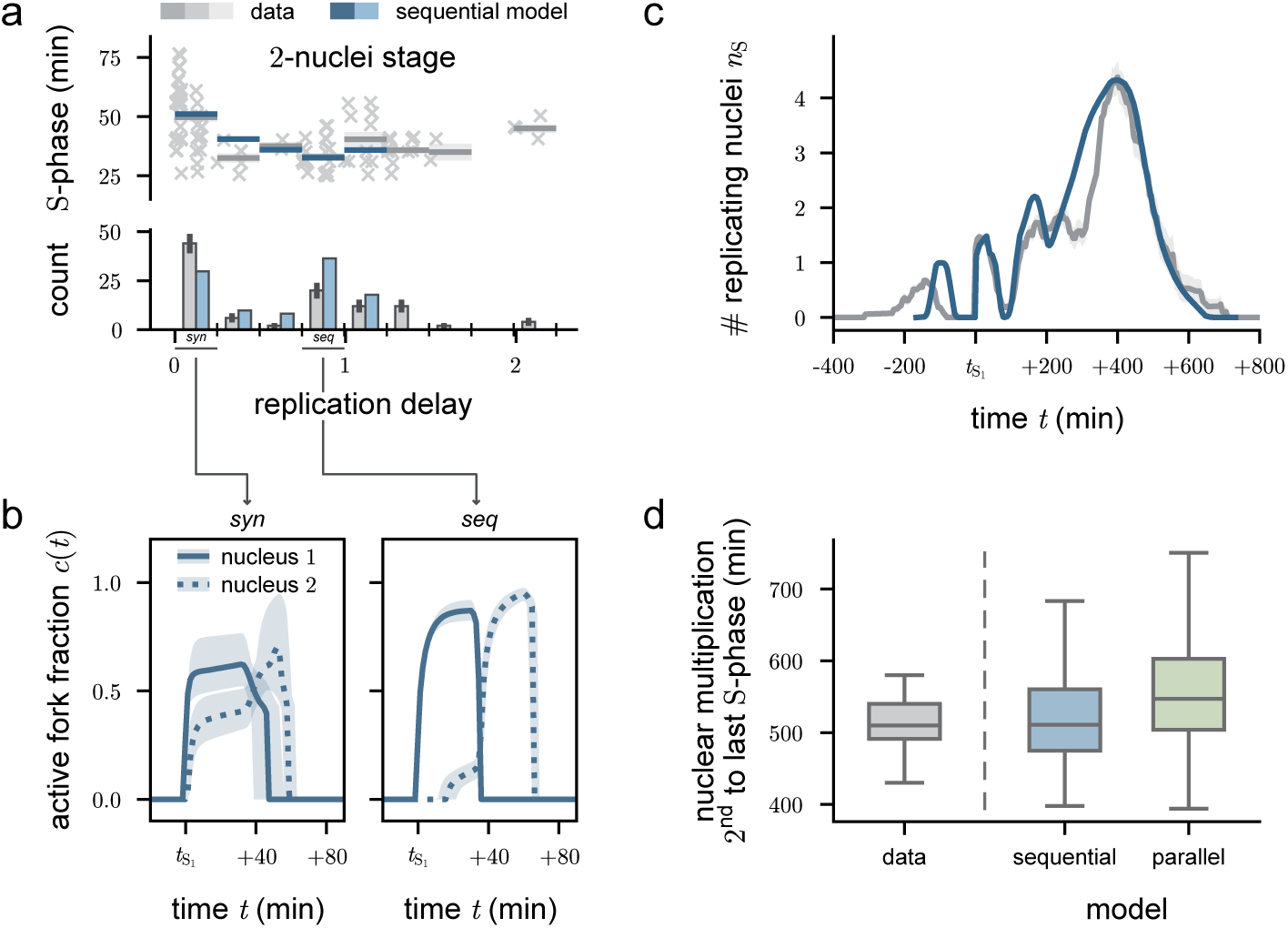
Model 3 captures nuclear multiplication dynamics in *P. falciparum* schizonts. (a) S-phase duration versus replication delay, as in Fig. 2d. Gray: replotted data, blue: best-fit model 3 in sequential mode, reproduces the prolonged simultaneous S-phases and the depletion of intermediate delays. (b) Fraction of active resource-fork complexes *c* at the 2-nuclei stage. Synchronous sisters (left) share resource and replicate slowly; delayed sisters (right) queue for resource and then replicate quickly. Lines and shading: simulation median and interquartile range, respectively. (c) Temporal profiles of the number of replicating nuclei. Desynchronization is similar in data and model 3. Colors as in (a). Profiles are aligned to the onset of the first S-phase in cells with two nuclei *t*_S1_). (d) Time to complete nuclear multiplication, from start of S_1_ to end of last S-Phase. In a-d, 10^5^ realizations were simulated, stopping at 24 nuclei. *τ* = 102 min, *τ*_S_∗ = 28 min, *σ*_D_∗ = 10 min, *ξ* ∼ Uniform(1.2, 2.3), *c*_min_ = 0.2. (*k_u_, k_b_*) = (9.8 · 10^−5^, 0.49) min^−1^ and (0.98, 4900) min^−1^ for best-fit (sequential mode) and parallel mode, respectively.

Furthermore, although model 3 was not parameterized using the observed correlations between the cycle phases of closely related nuclei, it successfully reproduced their correlation pattern, including cousins (Fig. S3b).

To more thoroughly test model 3, we assessed whether it can reproduce another feature of *P. falciparum* nuclear multiplication, the rapid desynchronization in the nuclear population over time. Using high-resolution time-lapse microscopy, we recorded DNA replication activity in *P. falciparum* schizonts. Fig. 6c shows the number of concurrent S-phases over time averaged over N = 49 parasites (Fig. 6c, gray line). A subset of DNA replications at the two-nuclei stage were synchronous, with a maximum average number of ≈ 1.4 simultaneous S-phases occurring in the peak after t_S_∗_1_ . This agrees with the partial synchrony already shown in Figs. 2d and 6a. Later on, a DNA replication gap with most nuclei in D^∗^-phase is observed after the 2-nuclei stage (t ≃ 100min) but no distinct gaps were detected after the 4-nuclei stage (t ≃ 200min). We then generated corresponding profiles employing model 3 with the parameter sets obtained above based on only 2-nuclei stage data. We found that when resource release is slow, gaps with a majority of nuclei in D^∗^-phase extend over several nuclear cycles, showing much slower desynchronization than observed experimentally (Fig. S4f). However, for slow resource release, e.g., in sequential mode, a fraction of nuclei are queued for replication in each generation. This successfully reproduces the rapid desynchronization from the 2-nuclei stage onward, including the loss of the gaps after 4 nuclei (Fig. 6d and S4). Thus, using the same sequential-mode parameters that explain prolonged simultaneous S-phases in early sister nuclei, model 3 also recapitulates the later desynchronizing effect of a limiting resource pool.

We conducted an additional set of experiments using a different microscope with higher resolution (Zeiss LSM900 equipped with an Airyscan 2 detector, see Methods). We found that now, nuclear cycles were consistently slower by about 33% (Fig. S5a). The overall slow-down is likely due to increased phototoxic effects when using the Zeiss microscope compared to the spinning disk PerkinElmer UltraVIEW VoX microscope used in the initial set of experiments. The new data again showed a slow-down of simultaneous sister S-phases as well as the bimodal distribution of S-phase delays, but fewer events with a replication delay around 1 were detected (Fig. S5b). Nuclei now also remained more synchronous for the entire duration of schizogony (Fig. S5c).

Perhaps surprisingly, these observations are, in fact, compatible with model 3, when it operates under conditions where the nuclear phases are slowed down, but resource production is unchanged. More precisely, when lengthening average S^∗^- and D^∗^-phases by 33% while keeping the time course r(t) and all other sequential-mode parameters unchanged, model 3 again matches the 2-nuclei stage statistics, the correlation pattern between the cycle phases and the slowed desynchronization of the repeat experiment (Fig. S5b-d). The increased synchrony arises in the model as follows. Because resource abundance is effectively increased when nuclei replicate slower, competition for resource is less severe in these conditions. With more resource, the lagging sister at the 2-nuclei stage can then begin S-phase sooner, and later stages desynchronize more slowly due to reduced queuing.

Taken together, our data support the following interpretation: When *P. falciparum* terminates the trophozoite stage and enters the schizont stage, the replication machinery (resource) is limiting for DNA replication. In good growth conditions, resource remains limiting for the duration of nuclear multiplication despite continued production. Sequential sharing of the limited resource, detectable as rapid desynchronization of nuclear phases, then ensures optimal utilization of replication machinery. In less favorable growth conditions, nuclear multiplication may be slowed down, e.g., by limited nutrient availability. Then the shared resource may not be limiting anymore, resulting in nuclear cycles that remain synchronous much longer. In these conditions, sequential vs. parallel resource sharing is not phenotypically different, and no speed benefits can be obtained.

## DISCUSSION

In this study, we have addressed the open questions of how asynchronous nuclear multiplication arises in *P. falciparum* schizonts, and what benefits it may have. After comparing statistical and biophysical models with imaging data, we conclude that *P. falciparum* has evolved resource sharing between nuclei in a common cytoplasm as a way to achieve efficient proliferation. The experimentally supported model features a limiting pool of a (co-)factors of DNA replication (resource) shared among nuclei through reversible loading on DNA and translocation. The model successfully reproduces both the slowdown of overlapping S-phases and the rapid desynchronization of nuclear cycles previously observed in *P. falciparum* [7, 13]. When the resource residence time at DNA replication forks is long compared to S^∗^-phase, then replicating nuclei sequester resource, enqueuing new S^∗^-phase nuclei for DNA replication. A central finding is that this sequential mode of DNA replication can speed up nuclear multiplication by shuttling replication resource between nuclei, increasing the fraction of time it is in use. Our results suggest that asynchronous multiplication of nuclei in *P. falciparum* is not a sign of autonomy, as in fungal hyphae [42], and argue against an entirely local control of *Plasmodium* nuclear cycles [8]. Rather, asynchrony appears as a manifestation of a physical coupling of nuclei via competition for resource.

While the molecular identity of resource R in the model is unspecified, our available data are consistent with R being a cofactor immediately involved in DNA replication, potentially identical to PCNA1 [13]. S^∗^-phases in *P. falciparum* also depend on *Pf* CRK4, a kinase that regulates a set of proteins involved in replication origin firing [6, 40]. In the context of the model, activity of *Pf* CRK4 may indicate or instruct S^∗^-phase entry, while R controls processive replication (S-phase). In addition, how nuclear multiplication is embedded in the overarching rythmicity of *Plasmodium* development (a multiple of 24 hours depending on the species [43]), is not addressed in the model and remains an interesting question for future research.

Apart from competition for the DNA replication cofactor R, other coupling mechanisms could also exist, but the most plausible alternatives are ruled out by the experimental findings. First, coupling could arise through a pool of a nutrient-like resource H which is full initially and then successively depleted as nuclei multiply (unlike R in our model), setting the final size of the system, very similar to limiting pool mechanisms for organelle growth [26]. However, H limitation would set in only after a certain number of nuclei have been produced, failing to generate the correlations between early nuclei seen in Fig. 2b. Second, DNA replication also depends on various consumable but renewable resources N, such as histone proteins and nucleotides. If running S^∗^-phases exhaust the current N pool, then N becomes limiting. This would slow DNA replication overall, but it would not delay S-phase entry in lagging sister nuclei, as observed. Last, the localization of R to nuclei could be driven by active nuclear import instead of DNA association. Although feasible, this mechanism would require cycle-dependent regulation of nucleocytoplasmic transport activity in addition to resource-fork binding kinetics, and including this in a model appears not warranted given the available data.

Sequential DNA replication speeds up nuclear multiplication, but does maximal proliferation in RBCs also maximize fitness of *P. falciparum*? Assuming that faster nuclear multiplication leads a higher merozoite number and/or a shorter overall schizogony, it will increase the rate of proliferation of the parasite in the host’s blood. Overly fast proliferation of pathogens has been argued to imply rapid host mortality, which would reduce transmission and therefore pathogen fitness [44–46]. However, for malaria with an overall sub-1% mortality and a substantial fraction of mild or asymptomatic infections [47], this so-called virulence-transmission trade-off does not appear relevant. Furthermore, at later stages of persistent disease, adaptive host immunity curtails RBC reinfection by merozoites so that maximizing merozoite production rate becomes crucial for continued transmission.

*P. falciparum* has evolved another trait that strongly suggests a selective pressure for speedy schizogony. Because infected RBCs have altered mechanical properties, schizonts are at risk of being eliminated when passing with the bloodstream through the mechanical RBC quality control system in the spleen. To counter the risk, parasites induce a system of adhesive knobs on the RBC surface [48] that increase the time between passages through the interendothelial slits in the spleen beyond the normal interval of around 5 hours [49]. In combination with adhesion, rapid nuclear proliferation will then increase the chances of completing the proliferative cycle before being detected by the RBC quality control.

The emerging picture is that *P. falciparum* is under pressure for rapid proliferation in an environment that limits resources for growth, as evidenced, e.g., by mechanisms to increase RBC permeability to nutrients [50]. This creates the fundamental challenge of allocating the limited resources available in a single RBC in order to maximize viable offspring in the limited time window between splenic passages. In particular, economizing on DNA replication machinery may have important advantages: First, *P. falciparum* can divert available supplies toward other purposes. Second, initiating multiplication when resource is not yet abundant means *P. falciparum* can proceed earlier. Both effects should reduce the total time to complete schizogony. To probe for signatures of maximized *P. falciparum* nuclear multiplication, we compared simulations of model 3 using the best-fit, sequential, parameter set, to simulations with unchanged parameters except that k*_u_*, k*_b_* were increased into the parallel replication regime while fixing the dissociation constant k*_b_*/k*_u_*. We found that when DNA was replicated in parallel, nuclear multiplication was slower by 40min ≃ 8% (Fig. 6d). This argues that the DNA binding kinetics of the replication machinery in *P. falciparum* may in fact be optimized for maximally efficient proliferation within RBCs.

Our work shows that resource sharing is not necessarily an impairment that has to be overcome in a quickly growing population, but can be a mechanism to organize growth for maximum efficiency, which could be highly relevant also for other developing cells or organisms. While bacterial communities are well-known to secrete and share enzymes, it is unclear if these can intermittently localize to different bacterial cells. Multinucleated eukaryotic cells with nuclei in close proximity could share the DNA replication machinery between nuclei by diffusion, making them candidates for efficient resource sharing. These occur during reproduction by multiple fission in protozoa, in particular Apicomplexa (of which *Plasmodium* schizogony is an example), but are also seen in unicellular algae. Fungal hyphae are another ubiquitous example of multinucleated cells. Their nuclei are separated spatially by several µm, which hinders diffusive sharing of replication machinery. It is an interesting open question whether they instead exploit periodic cytoplasmic flows for resource sharing.

In conclusion, we have found that the observed asynchronous nuclear cycles during *P. falciparum* schizogony can be explained by competition for a shared replication resource, expanding the regulatory repertoire assigned to limiting pool mechanisms [26, 27]. Competitive sharing allows *P. falciparum* to use replication machinery more efficiently, speeding proliferation under resource limitation, and increasing fitness in the hostile environment of the human host.

## Supporting information

Mathematical details of the nuclear multiplication models

## METHODS

### P. falciparum cell culture

The *P. falciparum* strain 3D7 is laboratory-adapted and is derived from the isolate NF54 by limiting dilution. NF54 originated from a case of airport malaria in the Netherlands [51]. *P. falciparum* 3D7 was cultivated in fresh O, Rh+ erythrocytes at 2 − 4% hematocrit in supplemented RPMI 1640 medium (with 0.2 mM hypoxanthine (CCPro), 25 mM HEPES, pH 7.4 (Merck), and 12.5 µg/ml gentamicin (Carl Roth), as well as 0.5% AlbuMAX II (Gibco)), at 37^◦^C in 90% relative humidity, 5% O2, and 3% CO2 [52]. Routine synchronizations were performed by 5% sorbitol treatment as previously described [53].

### P. falciparum cell lines

The PCNA1::GFP and NLS::mCherry expressing *P. falciparum* 3D7 line was previously generated [13]. To detect DNA replication events, PCNA1::GFP was expressed from the episomal plasmid pARL PCNA1-GS-eGFP under the control of the *P. falciparum* CRT promoter and the *P. berghei* DHFR/TS terminator.

*P. falciparum* PCNA1 is connected to eGFP via a 2 × GGGGS-linker. Additionally, this plasmid contains a WR99210 resistance cassette [54]. To visualize the nucleoplasm and detect nuclear division events, mCherry coupled to 3 nuclear localization sequences (NLS) was expressed under the control of the *P. falciparum* hsp86 promoter and the *P. berghei* DHFR/TS terminator. Furthermore, this plasmid contains a blasticidin (BSD) resistance cassette.

### Live-cell imaging

For live-cell imaging, we followed previously published protocols with minor modifications [13, 55]. Sterile glass bottom 8-well dishes (ibidi GmbH) were coated with 5 mg/ml Concanavalin A (Merck) and rinsed with PBS. To generate a monolayer of cells, approximately 500 µl of resuspended parasite culture were washed twice with supplemented RPMI without AlbuMAX II and left to settle on the dish for 10 min at 37^◦^C before unattached cells were washed off using supplemented RPMI without AlbuMAX II. Cells were left to recover at standard culturing conditions in supplemented RPMI with AlbuMAX II for at least 8 h before media was exchanged to the phenol red-free imaging medium (RPMI 1640 L-Glutamine, PAN-Biotech, 0.5% AlbuMAX II, 0.2 mM Hypoxanthine, 25 mM HEPES pH 7.3, 12.5 µg/ml gentamicin), which either had been equilibrated to incubator gas conditions for at least 6 h before sealing of the imaging dish or imaging was conducted in a gassed incubation chamber. Unless otherwise noted, long-term live-cell imaging was carried out on a PerkinElmer UltraVIEW VoX microscope equipped with Yokogawa CSU-X1 spinning disk head and Nikon TiE microscope body. An Apo TIRF 60x/1.49 N.A. oil immersion objective and Hamamatsu C9100-23B EM-CCD camera were used. Live-cell imaging was performed at 36.5^◦^C. Image were acquired at multiple positions using an automated stage and the Perfect Focus System (PFS) for focus stabilization with a time-resolution of 5 min/stack. Multi-channel images were acquired sequentially using solid state lasers with excitation at 488 nm and 561 nm and matching emission filters in addition to differential interference contrast (DIC) images; 8 µm stacks were acquired with a z-spacing of 500 nm.

For an independent data set, we used point laser scanning confocal microscopy, performed on a Zeiss LSM900 microscope equipped with an Airyscan 2 detector using Plan-Apochromat 63x/1,4 oil immersion objective. Live-cell imaging was performed at 37^◦^C. The images were acquired using the Airyscan detector in SR mode at multiple positions using an automated stage and the Definite Focus module for focus stabilization with a time-resolution of 5 min/stack for up to 18 h. Multichannel images were acquired sequentially using 488 nm and 561 nm diode lasers for eGFP and mCherry imaging, respectively. Emission detection was configured using variable dichroic mirrors to be 490 − 570 for eGFP and 570 − 650 for mCherry detection. The Airyscan detector was used with the gain adjusted between 700 and 900 V, offset was not adjusted (0%). Brightfield images were obtained from a transmitted light PMT detector, with the gain adjusted between 300 and 500 V. The sampling was Nyquist-optimized in xy axis (approx. 50 nm), and 600 µm in z axis, bidirectionally with pixel dwell time around 0.7 µs. Subsequently, the ZEN Blue 3.1 software was used for 3D Airyscan processing with automatically determined default Airyscan Filtering (AF) strength.

### Analysis of live-cell imaging data

The initial processing and analysis of the imaging data was done with Fiji [56]. For further analysis, we used Microsoft Excel. Multiposition images were manually inspected for dead or abnormal parasites using the DIC or brightfield channel. Individual data were saved as either 200 × 200 or 300 × 300-pixel TIFF files containing original channel, time and z-slice information. The time-lapse images were stabilized in xy as previously described [13]. In brief, either using the Register Virtual Stack Slices Plugin in Fiji, we registered for each parasite a time-lapsed reference z-slice of the brightfield channel via the translation mode (no deformation) at standard parameters. The transformation matrices were saved and applied to all other z-slices and channels using the Transform Virtual Stack Slices Plugin in Fiji. Alternatively, we used the Fiji ‘Correct 3D Drift’ plug-in [57], which corrects for movement in the xy plane within a specified maximum pixel shift, using a reference channel (here, brightfield).

The fluorescence intensity over time was previously published and re-analyzed for Fig. S3a [13]. Representative PCNA1::GFP expressing parasites were selected where both the onset of the first S-phase and egress were clearly visible. Raw integrated density was measured in the GFP channel of average intensity z-projections and normalized to maximal and minimal values per cell. Data from individual cells were aligned by the timepoint of parasite egress. We determined the dynamics of the nuclear cycle phases using our nuclear cycle sensor line and as previously described [13, 40]. In brief, S-phase was defined as the time interval between the onset and the end of visible PCNA1::GFP accumulation in a given nucleus. The interval between the end of the visible PCNA1::GFP accumulation and the ensuing accumulation was defined as division (D-) phase.

To quantify the number of nuclei in a given *P. falciparum* schizont (Fig. S2), we manually counted the number of mCherry-positive structures in each cell over time. Similarly, to determine the number of nuclei in S-phase at any given time point (Fig. 6c), we counted the number of mCherry-positive structures that also exhibited accumulated PCNA1::GFP signal within the same cell. For image visualization and quantification of nuclei and nuclei in S-phase, we used the Imaris software (Oxford Instruments), versions 10.1 and 10.2.

## ACKNOWLEDGEMENTS

The authors are grateful to DKFZ core funding, to the Infectious Diseases Imaging Platform (www.idip-heidelberg.org), and the Plasmodium database PlasmoDB (www.plasmodb.org), which facilitated this work. M.G. is supported by the Health + Life Science Alliance Heidelberg Mannheim, receiving state funding approved by the State Parliament of Baden-Württemberg. U.S., M.G. and T.H. received support from the German Research Foundation (Deutsche Forschungsgemeinschaft, DFG) through SFB 1129, project number 240245660, subprojects TP4, TP18 and TP11, respectively.

## Author contributions

Conceptualization, P. B., M. G. and N. B. B.; Methodology, P. B., M. G. and N. B. B.; Formal Analysis, P. B.; Investigation, P. B., A. K. and S. K.; Writing – Original Draft, P. B., A. K., S. K., M. G. and N. B. B.; Writing – Review & Editing, P. B., T. H., U. S. S., M. G. and N. B. B.; Supervision, M. G. and N. B. B.; Funding Acquisition, T. H., U. S. S. and M. G.

