## Supplementary material for "Competitive resource allocation drives asynchronous and rapid nuclear multiplication in the malaria parasite": Mathematical details of the nuclear multiplication models

Patrick Binder,[1,2] Aiste Kudulyte,[3] Severina Klaus,[3] Thomas Höfer,[1]
Ulrich S. Schwarz,[2] Markus Ganter,[3,*] and Nils B. Becker[1,†]

[1] *Theoretical Systems Biology, German Cancer Research Center (DKFZ), Heidelberg, Germany*

[2] *Institute for Theoretical Physics and BioQuant, Heidelberg University, Heidelberg, Germany*

[3] *Center for Infectious Diseases - Parasitology, Medical Faculty, Heidelberg University, Heidelberg, Germany*

(Dated: August 7, 2025)

### I. CONSTRUCTION AND CALIBRATION OF MODEL 2: BIFURCATING AUTOREGRESSIVE MODEL

In order to test if the temporal features of the nuclear cycle phases in *P. falciparum* can be explained by simple local inheritance, we introduce model 2. Model 2 is constructed as a bifurcating autoregressive process that incorporates the local correlations between the D- and S-phases existing between mother and both daughter nuclei. These correlations define a joint multivariate Gaussian distribution for mother and daughter phases with mean vector $\boldsymbol{\mu}$ and covariance matrix $\boldsymbol{\Sigma}$,

$$p(\mathbf{x}) = \frac{1}{\sqrt{\det(2\pi\boldsymbol{\Sigma})}} \exp\left[-\frac{1}{2}(\mathbf{x}-\boldsymbol{\mu})^{\intercal}\boldsymbol{\Sigma}^{-1}(\mathbf{x}-\boldsymbol{\mu})\right]. \tag{1}$$

Simulation of the branching process forward in time then involves drawing the daughter D- and S-phases conditioned on the given mother D- and S-phases, in agreement with Eq. 1. To obtain the required conditional distribution, we partition the multivariate Gaussian variable $\mathbf{x}$, its mean and covariance as

$$\mathbf{x} = \begin{bmatrix} \mathbf{x}_m \\ \mathbf{x}_d \end{bmatrix}, \quad \boldsymbol{\mu} = \begin{bmatrix} \boldsymbol{\mu}_m \\ \boldsymbol{\mu}_d \end{bmatrix} \text{ and } \quad \boldsymbol{\Sigma} = \begin{bmatrix} \boldsymbol{\Sigma}_{mm} & \boldsymbol{\Sigma}_{md} \\ \boldsymbol{\Sigma}_{dm} & \boldsymbol{\Sigma}_{dd} \end{bmatrix}, \tag{2}$$

where subscripts $\cdot_m$ and $\cdot_d$ correspond to the phases of the mother and to those of both daughters, respectively. It's well-known that the conditional distribution of $\mathbf{x}_d$ given $\mathbf{x}_m = \overline{\mathbf{x}}_m$ is also Gaussian, with adjusted mean $\overline{\boldsymbol{\mu}}_d$ and adjusted


*

†

covariance matrix $\overline{\mathbf{\Sigma}}_{dd}$, i.e.

$$\overline{\boldsymbol{\mu}}_d = \boldsymbol{\mu}_d + \mathbf{\Sigma}_{dm}\mathbf{\Sigma}_{mm}^{-1}(\overline{\mathbf{x}}_m - \boldsymbol{\mu}_m)\,, \tag{3a}$$

$$\overline{\mathbf{\Sigma}}_{dd} = \mathbf{\Sigma}_{dd} - \mathbf{\Sigma}_{dm}\mathbf{\Sigma}_{mm}^{-1}\mathbf{\Sigma}_{md}\,. \tag{3b}$$

To forward simulate a standardized Gaussian version of a nuclear population, we begin from the founder nucleus' S-phase, as the preceding initial D-phase is not available from experiment. Thus, for the initial generation only, $\mathbf{x}_m$ reduces to a one-dimensional standard Gaussian variable, and in subsequent generations, $\mathbf{x}_m$ is two-dimensional (covering both D- and S-phases). $\mathbf{x}_d$ is a four-dimensional standard Gaussian for the daughters' D- and S-phases.

We next map this Gaussian bifurcating process onto the correct empirical phase distributions. Specifically, the empirical mother-phases, denoted by $\mathbf{x}_m^{\text{ex}}$, are transformed into standard Gaussian variables $\mathbf{x}_m$ using the nonlinear mapping

$$\mathbf{x}_m = g\big(\mathbf{x}_m^{\text{ex}}\big) = c_{\text{gauss}}^{-1}\Big(c_{\text{ex}}\big(\mathbf{x}_m^{\text{ex}}\big)\Big)\,, \tag{4}$$

where $c_{\text{gauss}}$ denotes the cumulative distribution function (CDF) of the standard Gaussian distribution and $c_{\text{ex}}$ is the CDF corresponding to the empirical data. Conversely, simulation data matching the empirical phase distributions can be generated from Gaussian samples by the inverse mapping

$$\mathbf{x}_m^{\text{ex}} = g^{-1}(\mathbf{x}_m). \tag{5}$$

Thus, to simulate nuclear populations in model 2, we first sample a standard Gaussian for the initial S-phase, then successively apply Eq. 3 to produce a Gaussian correlated tree, and finally transform onto the correct marginal distributions using Eq. 5.

To parametrize model 2, we first study the empirical phase distributions. Specifically, we investigate whether empirical distributions from related cells can be pooled to construct more robust cumulative distribution functions $c_{\mathrm{ex}}$ for the S- and D-phases.

As reported in [1], the very first S-phase is significantly prolonged compared to later S-phases. To account for this difference, model 2 uses this specific prolonged empirical distribution for the initial S-phase.

We next examine whether S-phases at the 2- and 4-nuclei stages follow a common distribution. Using the Kolmogorov-Smirnov (KS) test to compare each individual phase distribution to the S-phase durations pooled across the two stages confirms that S-phases can be treated as independent samples from a common distribution (Fig. S1a). Furthermore, later stages do not significantly deviate as data are scarce. Consequently, we model all S-phases starting from the 2-nuclei stage using the pooled distribution.

D-phases at the 2-nuclei stage are ordered such that the first sister nucleus always has a shorter D-phase than the second, i.e. $D_1 \leq D_2$, by convention. Indeed, $D_1$-phases follow a distribution shifted to shorter times compared to $D_2$. This effect may either simply be an artifact of our ordering convention, or there may exist an intrinsic asymmetry in the nuclear division process that by itself already generates a faster and a slower D-phase. To scrutinize this possibility, we constructed a symmetrized distribution by drawing 50 000 independent pairs of D-phases from the pooled empirical D-phase distribution, assigning the faster value to $D_1$ and the slower value to $D_2$. Using the two-sample KS test, we find that the so constructed $D_1$ and $D_2$-phase distributions are statistically indistinguishable from the corresponding empirical distributions (Fig. S1a). We conclude that the data do not support asymmetric division, and model the 2-nuclei stage D-phases as symmetric and independent, using the pooled empirical distribution. A similar analysis for the 4-nuclei stage revealed that the D-phases of all four sister and cousin nuclei can equally be described as independent samples from a common distribution (Fig. S1a). Therefore, we model the D-phases at the 4-nuclei stage using the pooled distribution of the empirical data from $D_{11}$, $D_{12}$, $D_{21}$, and $D_{22}$. Because no later D-phases were observed, we continue to use this pooled distribution to model all subsequent D-phases.

The final model 2 comprises four empirical phase distributions in total. One S-phase and one D-phase distribution describe the initial phases, while two more distributions, pooled from later data, describe all subsequent S-and D-phases, respectively (Fig. S1b). To avoid introducing a censoring bias in pooling, we include a nuclear phase only if the corresponding phase was also observed in the sister nucleus.

We next discuss our procedure for extracting correlations from data. First, we transform the empirical data into standard Gaussian variables using Eq. 4. The Gaussian rank correlations then equal the moment correlation coefficients between these transformed variables. Because the imposed ordering of sister D-phases ($D_1 \leq D_2$) would introduce a bias towards positive correlation, and in accordance with our independent symmetric model, we computed the sister correlation of the D-phases with shuffled pairs. Based on the estimated Gaussian rank correlations, we then constructed a parsimonious covariance matrix that retains the significant mother-daughter and sister correlations but sets all non-significant correlations to zero. Bootstrapping confirmed that the common covariance Gaussian rank correlation matrix $\Sigma_{\text{Gaussian}}$ is sufficient to capture the significant correlations across the different nuclear stages (Fig. S1c). By defining the phase vector $\Phi = \left[\text{D S D}_a \text{ D}_b \text{ S}_a \text{ S}_b\right]^\mathsf{T}$ ($\cdot_a$ and $\cdot_b$ denote the two daughters), the Gaussian rank correlation matrix reads

$$\Sigma_{\text{Gaussian}} = \left\langle g(\Phi)g(\Phi)^{\mathsf{T}} \right\rangle = \left(\begin{array}{cc|cccc} 1 & 0 & 0 & 0 & 0 & 0 \\ 0 & 1 & 0 & 0 & 0.38 & 0.38 \\ \hline 0 & 0 & 1 & 0.45 & 0 & 0 \\ 0 & 0 & 0.45 & 1 & 0 & 0 \\ 0 & 0.38 & 0 & 0 & 1 & 0.79 \\ 0 & 0.38 & 0 & 0 & 0.79 & 1 \end{array}\right). \tag{6}$$

Note that since standard Gaussian variables have unit variance, the Gaussian rank correlation matrix is identical to the covariance matrix. Consequently, this matrix can be used directly for multivariate Gaussian sampling according to Eqs. 3.

### II. SEQUESTRATION OF RESOURCE

Sum Eq. 2 over $i$ to obtain at saturation

$$0 \simeq \dot{c} = k_b n_{\mathrm{S}^*} r^{\mathrm{free}} - (k_u + k_b r^{\mathrm{free}})(r - r^{\mathrm{free}}), \tag{7}$$

which yields a free fraction $r^{\mathrm{free}}/r = \frac{1}{n_{\mathrm{S}^*} - r} \frac{k_u}{k_b} + \mathcal{O}(\frac{k_u}{k_b})^2$. At high affinity, this is indeed $\ll 1$.

### III. CRITICAL AVAILABILITY

Because nuclei carry at least one genome, $\zeta \geq 1$ in Eq. 5 implies that $r \geq n \geq n_{\mathrm{S}^*}$, which allows full rate of replication in all nuclei. This bound is not sharp: Most of the time, some fraction of nuclei is in D$^*$-phase and does not compete for resource, so that a critical resource availability $< 1$ suffices for full speed replication. The critical value will decrease with increasing D*-to-S*-phase ratio.

### IV. POPULATION GROWTH RATE

To derive the population growth rate, consider a distribution of interdivision times $\tau$ with division time density $p(\tau)$ and survival function $S(a) = \mathbb{P}[\tau > a]$ valid for any newborn individual. After steady growth has been achieved, the average population $n(t)$ grows exponentially at (unknown) rate $\lambda$. The average total rate of divisions at time $t$ satisfies

$$\lambda n(t)dt = n(t)\int_0^t \mathbb{P}[\text{age } a \mid \text{alive at } t]\mathbb{P}[\text{division in } (t, t+dt) \mid \text{age } a \text{ at } t]da. \quad (8)$$

The division propensity conditioned on survival to age $a$ is given by

$$\mathbb{P}[\text{ division in } (t, t+dt) \mid \text{age } a \text{ at } t]/dt = -\dot{S}(a)/S(a). \quad (9)$$

The stationary age distribution can be obtained using Bayes' rule, by considering a base ensemble of all individuals born between 0 and $t$:

$$\begin{aligned}
&\mathbb{P}[\text{ age } a \mid \text{alive at } t]da = \\
&= \mathbb{P}[\text{ born in } (t-a-da, t-a) \mid \text{alive at } t] \\
&= \frac{\mathbb{P}[\text{ alive at } t \mid \text{born in } (t-a-da, t-a)]\mathbb{P}[\text{ born in } (t-a-da, t-a)]}{\mathbb{P}[\text{ alive at } t]} \\
&= \frac{S(a)\, e^{\lambda(t-a)}da/\int_0^t e^{\lambda(t-a')}da'}{\int_0^t S(a)\, e^{\lambda(t-a)}/\int_0^t e^{\lambda(t-a')}da'da} \\
&= \frac{S(a)e^{-\lambda a}da}{\int_0^t S(a')e^{-\lambda a'}da'}. \quad (10)
\end{aligned}$$

Here on the fourth line, we used the fact that births occur at a rate proportional to the exponentially growing population in the steady growth regime. Combining, Eq. 8 becomes

$$\begin{aligned}\lambda &= \frac{-\int_0^t \dot{S}(a)e^{-\lambda a}da}{\int_0^t S(a')e^{-\lambda a'}da'} \\ &= \frac{-\lambda\int_0^t S(a)e^{-\lambda a}da - [S(a)e^{-\lambda a}]_0^t}{\int_0^t S(a')e^{-\lambda a'}da'} \\ &\to -\lambda + \frac{1}{\int_0^\infty S(a')e^{-\lambda a'}da'},\end{aligned} \tag{11}$$

where we have integrated by parts and taken the limit $t\lambda \gg 1$. Solving Eq. 11 for the term appearing in the denominator yields $1/(2\lambda)$. Inserting this result and using $p(\tau) = -\dot{S}(\tau)$, the numerator becomes

$$\int_0^\infty p(\tau)e^{-\lambda\tau}d\tau = \frac{\lambda}{2\lambda} = \frac{1}{2}. \tag{12}$$

Substituting $p(\tau) = p(\tau_{\mathrm{D}^*})$ where $\tau = \tau_{\mathrm{S}^*} + \tau_{\mathrm{D}^*}$ for a constant $\tau_{\mathrm{S}^*}$, this is seen to imply Eq. 9.

### V. SIMULATION DETAILS FOR MODEL 3

To characterize the asymptotic growth regime of model 3, we simulated nuclear multiplication until the lineage tree contained $2^{15} - 1$ nuclei entering S*-phase, corresponding approximately to $2^{14}$ terminal nuclei. This simulation depth proved sufficient to reach and maintain the steady-growth regime, as evident from Fig. 4. To extract the steady growth rate $\lambda$, we determined the steady-state nuclear cycle duration $\tau_\infty$ as the time required for a nucleus to complete a full cycle (D*-phase followed by S*-phase) under steady-state conditions. (In the limit of deterministic

$\tau_{\mathrm{D}^*}$ and $\tau_{\mathrm{S}^*} + \tau_{\mathrm{D}^*} = 1$, the expression for $\tau_\infty$ reads $\tau_\infty = \min\left(\tau_{\mathrm{S}^*} \log 2/\zeta, 1\right)$.) We then fitted an exponential function to the total DNA content $g$ over approximately the final three cycles of the simulated data.

Similarly, the steady resource utilization $\eta$ was obtained as the average of $c/r$ across the final three cycles of simulations. The growth advantage $g_{\mathrm{seq}}/g_{\mathrm{par}}$ was read off at the end of the simulation, where it had converged sufficiently.

### VI. ADAPTING MODEL 3 TO *P. FALCIPARUM*

To match experimental data from time-lapse microscopy, we modified model 3 to represent the dynamics of *P. falciparum* nuclear multiplication more realistically. First, we rescaled the nuclear cycle duration in the model to match experimentally observed timescales. Because the initial phases are prolonged compared to later cycles [1], we used $\mathrm{S}_i + \mathrm{D}_{ij}$ as a reference for temporal mapping. Specifically, after simulating the model with the original parameters, we mapped the simulated S$^*$- and D$^*$-phases with $c_{\mathrm{min}} = 0.2$ to the corresponding S- and D-phases (see Fig. 3). We then rescaled time according to $t \to t/\tau_{\mathrm{rescale}}$, where the scaling factor is defined as $\tau_{\mathrm{rescale}} = \langle \mathrm{S}_i + \mathrm{D}_{ij} \rangle_{\mathrm{model}} / \langle \mathrm{S}_i + \mathrm{D}_{ij} \rangle_{\mathrm{ex}}$, with averages taken over all simulated and experimental cycles, respectively.

Next, we replaced Eq. 5 with a more realistic model for resource availability based on the measured time course of ectopically expressed PCNA1::GFP in *P. falciparum* [1] (Fig. S3a). The time course is approximated well by a saturating function with exponential rate $\gamma$ and scaling factor $\xi$:

$$r(t) = \xi\left(1 - e^{-\gamma t}\right) + r_0 e^{-\gamma t}. \tag{13}$$

Because the onset times of the second S-phases are not available for individual cells, we aligned all time traces to the egress event (merozoite release). We set the egress time to $t = 817$ min, corresponding to the experimentally observed average interval from the onset of the second S-phase to egress [1]. With this alignment, the onset of the second S-phase occurs approximately at $t = 0$, with the initial resource level $r_0$.

To infer the model parameters, we preprocessed the data through the following steps. First, we excluded non-responding cells, i.e., cells with almost no growth, indicated by a total fold-change $< 1.25$. Second, we subtracted a background signal defined as the initial value of the GFP signal from each trace, so that all traces start at a signal strength of 0. Third, we rescaled the traces so that their final plateau is given by our scaling parameter $\xi$, focusing on the change in signal. We then estimated parameter values $r_0 = 0.235\xi$ and $\gamma = 4.1 \cdot 10^{-3}$ min$^{-1}$. The choice of this saturating model is further supported by the observation that 3xNLS::mCherry signal in [1] follows a comparable saturating function as PCNA1::GFP.

[1] S. Klaus, P. Binder, J. Kim, M. Machado, C. Funaya, V. Schaaf, D. Klaschka, A. Kudulyte, M. Cyrklaff, V. Laketa, T. Höfer, J. Guizetti, N. B. Becker, F. Frischknecht, U. S. Schwarz, and M. Ganter, Science Advances **8**, eabj5362 (2022), https://www.science.org/doi/pdf/10.1126/sciadv.abj5362.

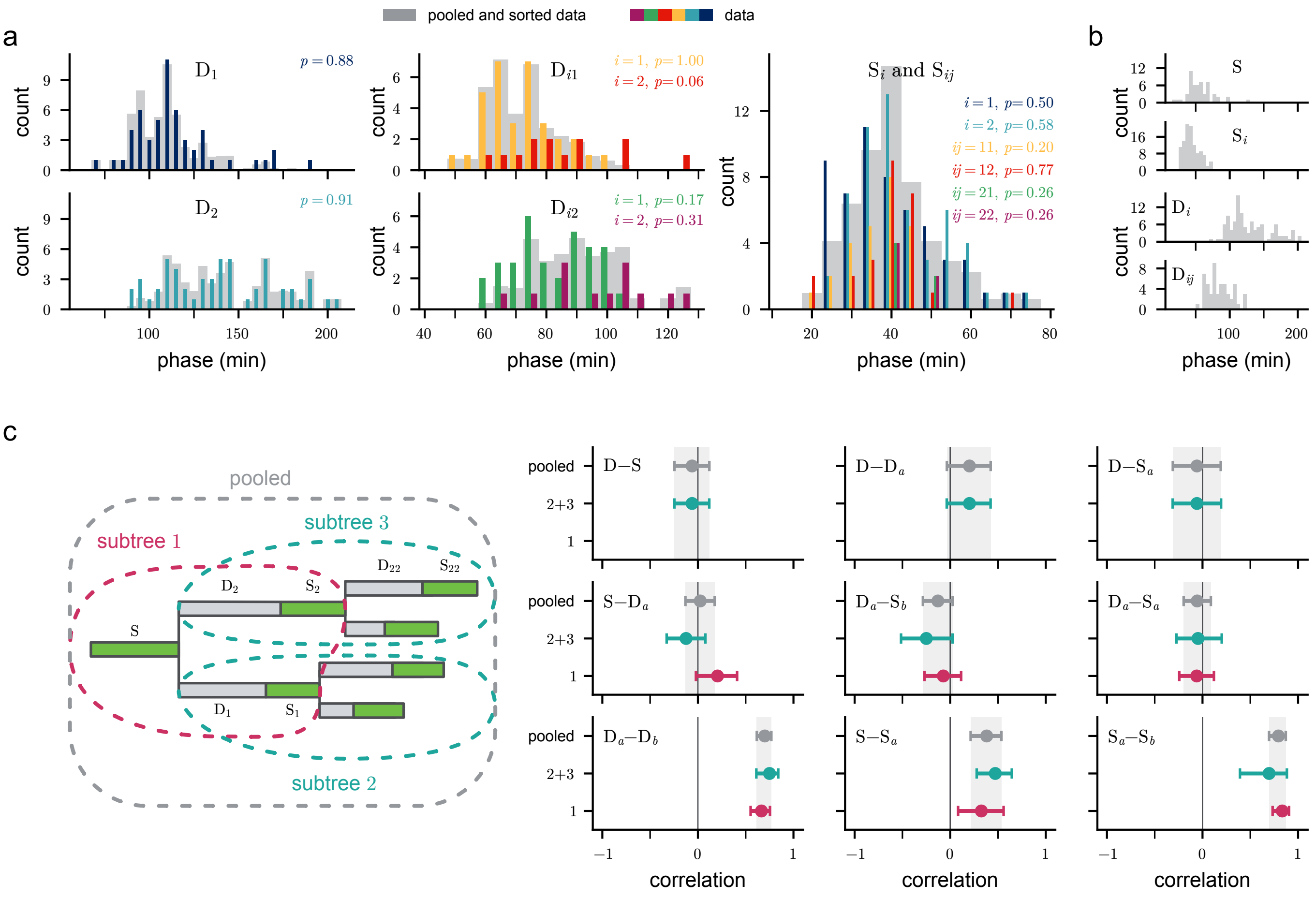


FIG. S1. Overview of model 2 construction from empirical phase distributions and correlations. (a) Distributions of S-phase durations ($S_i$, $S_{ij}$) and D-phase durations ($D_i$, $D_{ij}$) at the 2- and 4-nuclei stages, demonstrating that each phase type can be described by a single pooled distribution. $p$-values: two-sample Kolmogorov-Smirnov tests. (b) Pooled empirical distributions for S- and D-phases, which are used as input for model 2. (c) Left: Schematic of analyzed subtrees. Right: Gaussian rank correlations (mother-daughter and sister correlations) for all three subtrees, showing that a common correlation matrix accurately describes nuclear lineages.

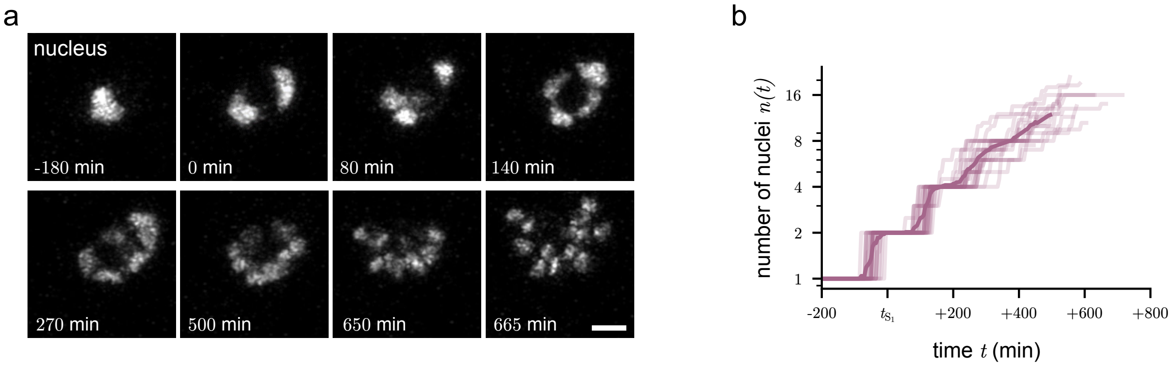


FIG. S2. *P. falciparum* nuclear multiplication dynamics. (a) Time-lapse microscopy of the reporter parasite expressing 3xNLS::mCherry using a Zeiss LSM900 equipped with an Airyscan 2 detector; shown are maximum intensity projections of the mCherry channel; scale bar, 2 µm. (b) Number of nuclei in a *P. falciparum* schizont over time. Initial nuclear multiplication of *P. falciparum* is close to exponential. Traces from $N = 24$ cells were aligned at the start of $S_1$: Light: single trace. Dark: mean. Counts are reliable up to $t \approx 400$ min and approximate afterwards. 4 traces where tracked until egress, the remaining are truncated.

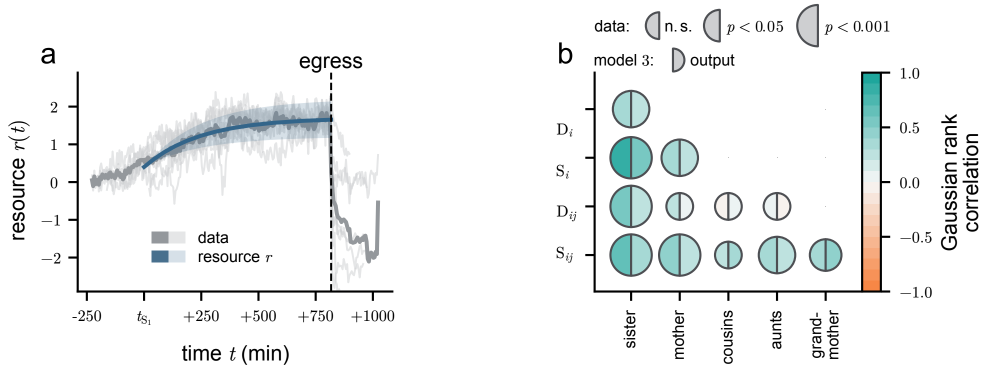


FIG. S3. Overview of resource dynamics and nuclear cycle correlations in model 3. (a) Parametrized resource growth in model 3 (blue) is inferred from shifted and rescaled total PCNA1-fluorescence signal (gray, see [1] Fig. S2C). PCNA1-fluorescence signal is rescaled such that each trace starts at a signal strength of 0 and ends at $\zeta$. All traces were aligned to the egress. Blue line and shaded area: mean and range of realizations due to variability in $\zeta$, where $\zeta$ is sampled uniformly. Light and dark gray: single trace and mean, respectively. (b) Correlation structure. Left-half disks: data. Right-half disks: model 3. Parameters as in Fig. 6.

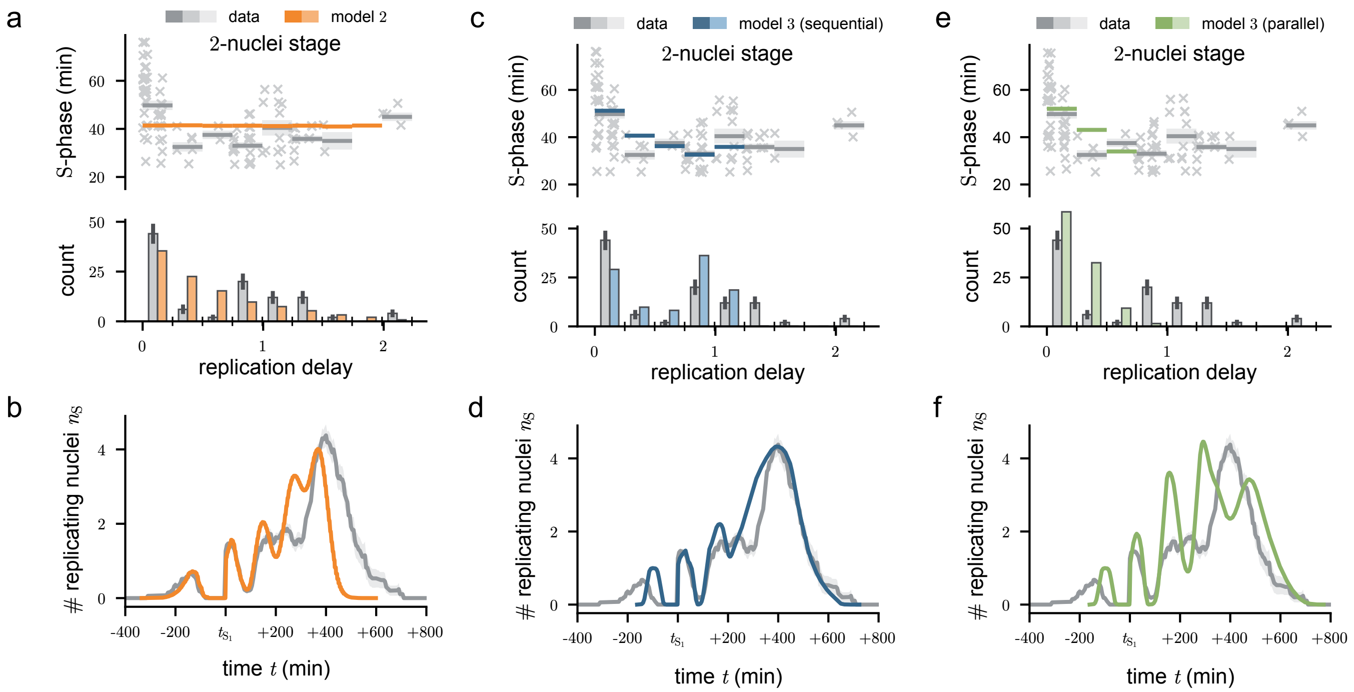


FIG. S4. Comparison of model 2 (inheritance-based, uncoupled nuclei) and model 3 (resource-sharing, coupled nuclei) in reproducing key features of nuclear multiplication dynamics. (ace) Duration of S-phase versus replication delay (cf. Fig. 2c). (bdf): Temporal profiles of the number of replicating nuclei (cf. Fig. 6c). Gray: experimental data. Orange (ab): model 2. Blue (cd): best-fit model 3 in sequential mode. Green (ef): model 3 in parallel mode. This comparison highlights that only model 3 with sequential replication can reproduce both the observed asynchrony and the correlation between replication timing and duration.

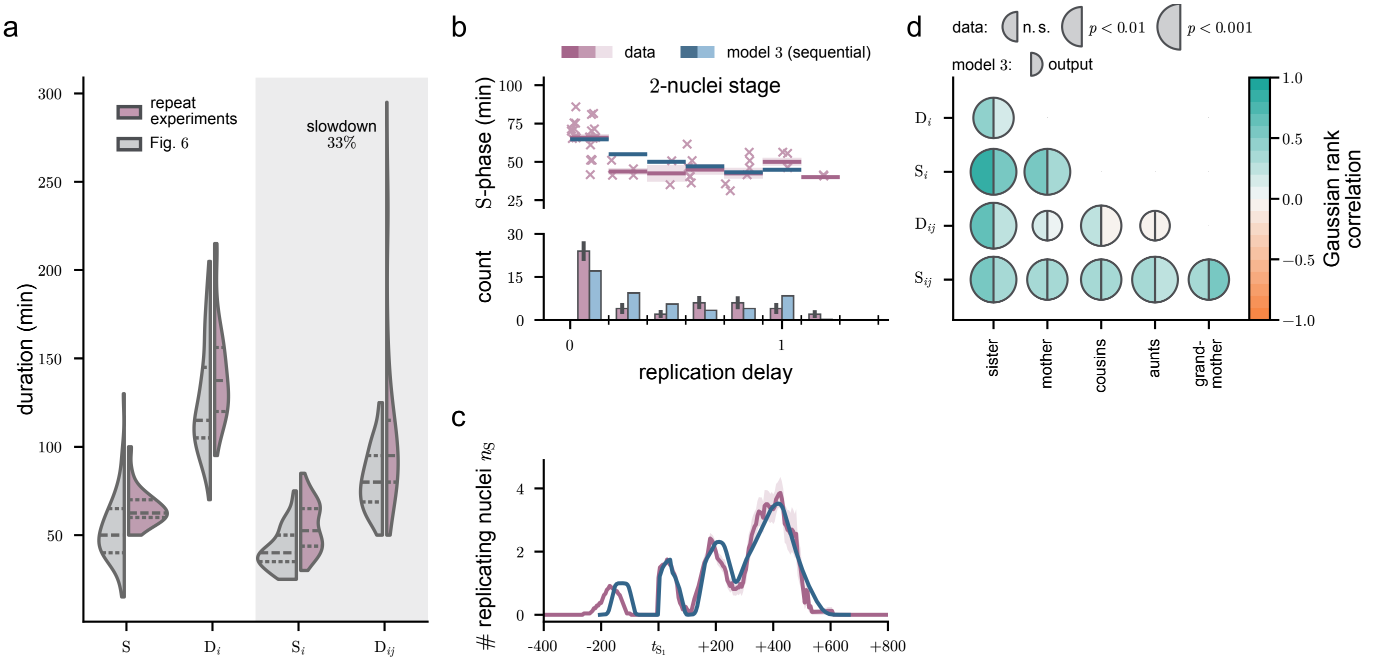


FIG. S5. Model 3 captures the nuclear multiplication dynamics of the repeat experiment in *P. falciparum* schizonts. (a) The initial two S- and D-phase durations are overall slower in the repeat experiment. Gray: data from main text; red: repeat experiment. As the very first phases are prolonged [1], we use $S_i$ and $D_{ij}$ (highlighted in light gray) to estimate the overall slowdown. (b) S-phase duration vs. delay. The sequential-mode model 3 (blue) captures prolonged and enriched simultaneous S-phases as well as depleted intermediate-delay S-phases. (c) The model successfully reproduce the observed relatively synchronous temporal profile of replicating nuclei. (d) Gaussian rank correlation structure as in Fig. 2c. In a-d, $10^5$ simulation realizations were run, each stopping at 16 nuclei. $\tau$ = 134 min, $\tau_{S*}$ = 37 min. All other parameters as in Fig. 6. In c, traces from $N = 24$ cells were aligned at the start of $S_1$, and averaged. 5 traces were tracked until egress; the remaining traces were followed up to approximately $t \approx 400$ min, after which data are incomplete.
